# Fast scanning high optical invariant two-photon microscopy for monitoring a large neural network activity with cellular resolution

**DOI:** 10.1101/2020.07.14.201699

**Authors:** Keisuke Ota, Yasuhiro Oisi, Takayuki Suzuki, Muneki Ikeda, Yoshiki Ito, Tsubasa Ito, Kenta Kobayashi, Midori Kobayashi, Maya Odagawa, Chie Matsubara, Yoshinori Kuroiwa, Masaru Horikoshi, Junya Matsushita, Hiroyuki Hioki, Masamichi Ohkura, Junichi Nakai, Masafumi Oizumi, Atsushi Miyawaki, Toru Aonishi, Takahiro Ode, Masanori Murayama

## Abstract

Fast and wide imaging with single-cell resolution, high signal-to-noise ratio and no optical aberration has the potential to open up new avenues of investigation in biology. However, this imaging is challenging because of the inevitable tradeoffs among those parameters. Here, we overcome the tradeoffs by combining a resonant scanning system, a large objective with low magnification and high numerical aperture, and highly sensitive large-aperture photodetectors. The result is a practically aberration-free, fast scanning high optical invariant two-photon microscopy (FASHIO-2PM) that enables calcium imaging from a large network composed of ∼16k neurons at 7.5 Hz in a 9 mm^2^ contiguous image plane including more than 10 sensory-motor and higher-order regions of the cerebral cortex in awake mice. Through a network analysis based on single-cell activities, we discover that the brain exhibits small-world-ness rather than scale-freeness. FASHIO-2PM will enable revealing biological dynamics by simultaneous monitoring of macroscopic activity and its composing elements.

## Introduction

If a phenomenon in a certain system (e.g., biology, social science, or physics) is observable, no matter how complex it appears to be, its mechanisms can be predicted by subdividing it into individual elements, but not vice versa. Given only the details of individual elements, the appearance of the phenomenon may not be predictable(*1*). The development of a method to simultaneously monitor a phenomenon and its elements will be a driving force for discoveries and open up new horizons in any field that benefits from observation. In neuroscience, the elementary information processing required for cognitive processes is thought to be executed within single brain regions, but emergent properties of the brain may require network activity involving multiple regions via long-/mid-range connections between neurons(*2, 3*). Thus, monitoring a large number of elements (i.e., single neurons) from multiple brain areas is necessary for a comprehensive understanding of brain functions, which is seen as one of the important challenges in this field. Most of the gains are currently being made via electrical high-density approaches(*4*) that make it difficult to identify or select the underlying cells or its geometric structures. Optical approaches can overcome this issue but quickly face intrinsic limitations.

To optically monitor a large number of neurons, one needs a microscope that has a wide FOV and a higher spatial resolution. However, these parameters are inversely related. To counteract this tradeoff, an imaging system requires a newly developed large objective with low magnification (Mag) and a high numerical aperture (NA). Recent efforts to achieve a wide-FOV two-photon (2P) excitation microscope take a strategy to increase the number of spatially separated small FOVs (each FOV being approximately 0.25 mm^2^) by employing this large objective(*5–7*). One of the systems, known as the Trepan-2P (Twin Region, Panoramic 2-photon) microscope(*5*), can also offer a contiguous wide-FOV imaging mode (i.e., without image stitching) because it has a high optical invariant (the product of the FOV and the NA; see the results for further details). This mode, however, significantly decreases the sampling rate (3.5 mm^2^ FOV for 0.1 Hz), resulting in a significant loss in temporal information in neuroscience. Another strategy that does not involve new large objective lenses but instead off-the-shelf components while increasing the optical invariant(*8, 9*) demonstrated ultrawide-FOV 2P imaging. Bumstead et al. achieved a 7 mm-diameter-FOV 2P microscope with a higher resolution in the lateral direction (1.7 μm). However, the axial resolution is limited by 28 μm and a significantly slow sampling rate because of the objective’s low performance based on the commercial objective, which is inadequate for single-cell imaging, and because a galvo-galvo scanning system is used, respectively. The ultrawide-FOV (6 mm in diameter) one-photon confocal microscope known as the Mesolens system(*10*), which also has a high optical invariant owing to the combination of a large objective lens and a galvo-galvo scanning system, achieves subcellular resolution with practically no aberration. However, the system may limit the imaging depth in biological tissues that have high light-scattering properties because it is designed for visible lasers(*11*). To the best of our knowledge, an optimized method that can permit fast imaging of large areas with single-cell resolution and no aberrations has yet to be fully examined. Nonetheless, previous efforts have made it clear that the strategy of maximizing optical invariant with high-performance large optics is a straightforward approach to realizing this microscope.

In this study, we maximized the optical invariant of a 2P microscope with a large-angled resonant scanning system, a newly developed well-designed (Strehl ratio, SR, ∼ 0.99 over the FOV) large objective (0.8 NA, 56 mm pupil diameter) and a large-aperture GaAsP photomultiplier (PMT, 14 mm square aperture). This combination is a new, untested approach for realizing fast, wide-and contiguous-FOV 2P imaging with single-cell resolution, a high signal-to-noise ratio (SNR) and practically no aberration across an entire FOV. Our microscope allowed us to monitor neural activity from ∼16000 cortical neurons in a contiguous 3 x 3 mm^2^ FOV at a 7.5 Hz sampling rate during animal behaviors. Using our microscope, we performed a functional network analysis based on a large number of single neurons and demonstrated the network properties of the brain.

## Results

Here, we describe the development of a fast scanning high optical invariant (0.6 for excitation and 1.2 for collection) two-photon microscopy (FASHIO-2PM) (see Table S1 for acquisition modes). Our microscope achieves concurrently the following six capabilities: 1) a contiguous wide FOV (3 x 3 mm^2^, 36x larger than that achieved with conventional ∼0.5 x 0.5 mm^2^ FOV microscopes(*12*); 2048 x 2048 pixels), 2) a high NA for single-cell resolution (optical resolution: x-y, 1.62 μm; z, 7.59 μm; see below), and 3) a fast frame rate (7.5 Hz) for monitoring neural activity.

### Roadmap for the FASHIO-2PM: Optical invariant

In an aberration-free and vignetting-less system, the optical invariant(*13, 14*), which is calculated with the height and angle of the chief and marginal rays, is conserved as a constant value in each system component. The optical invariant for the excitation light *I*_e_ of components ranging from the scanner plane to the image plane of a laser scanning 2P-microscope (LS2PM) with a non-descanned detector is as follows:

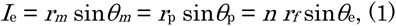

where *r_m_* and *θ_m_* are the beam radius and the scan angle at the mirror scanner, respectively, *r_p_* and *θ_p_* are the beam radius and the incident angle of collimated light at the pupil of the objective, respectively, *n* is the refractive index of the immersion between the objective and a specimen, *r_f_* is the FOV radius and *θ*_e_ is the angle of the cone of excitation light at the image plane (Fig. 1A). To avoid confusion, we define the plane at the rear of the objective near the tube lens as the rear aperture. Note that the last equation in Eq. (1) corresponds to the product of the NA and FOV radius. Importantly, the optical invariant for excitation *I_e_* is limited by the lowest optical invariant at one of the system’s components (i.e., the bottleneck component). Thus, increasing the lowest optical invariant at the bottleneck component will concurrently achieve a contiguous wide FOV and a high NA.

**Fig. 1.**
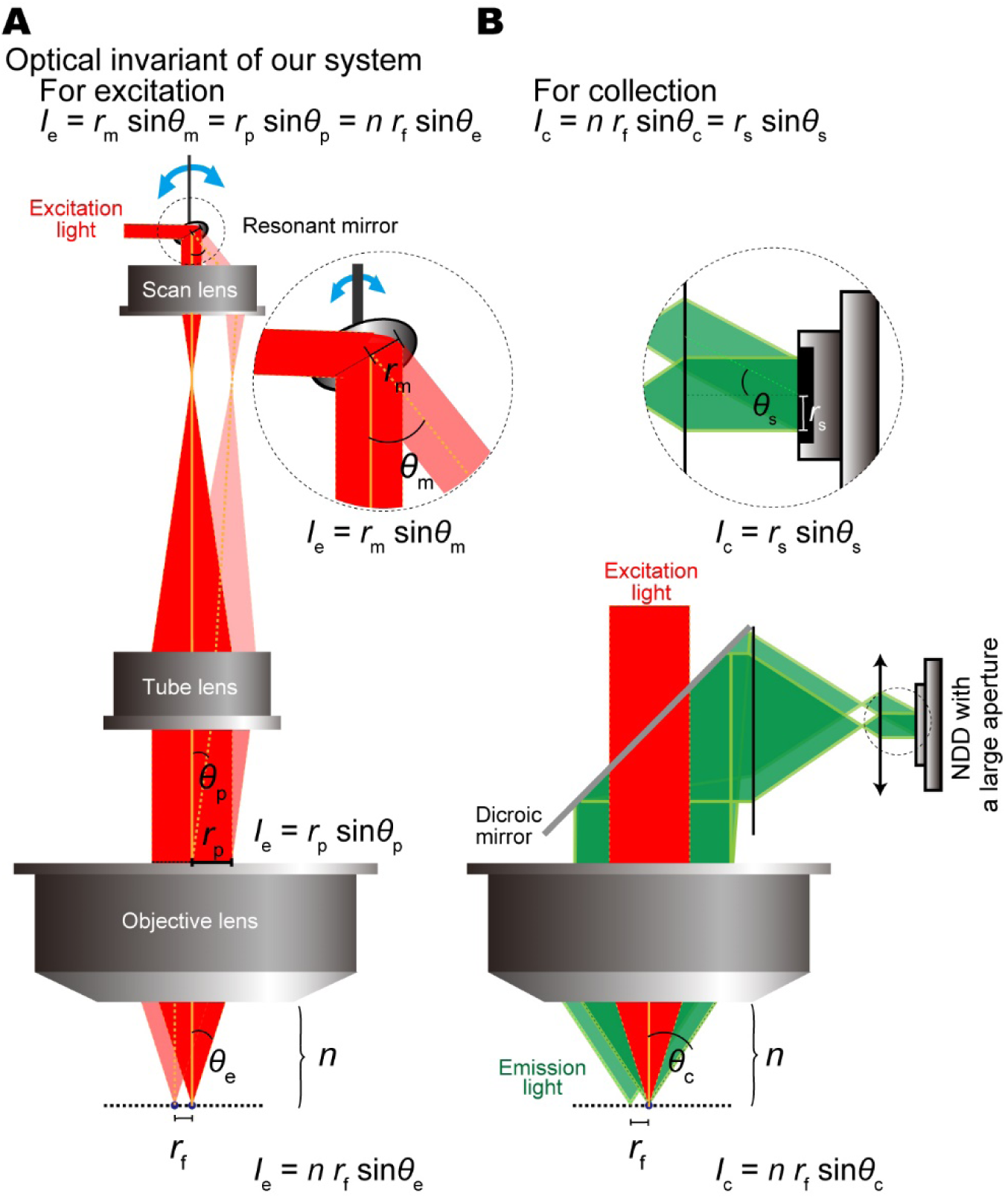
Optical invariant for the FASHIO-2PM. (**A**) Optical invariant for excitation light (*I*_e_) in an LS2PM. A collimated laser beam is scanned by a resonant mirror and projected to a pupil of the large objective lens through scan and tube lenses. The optical invariant at the object (i.e., specimen) side is equal to the invariant at the scanning mirror and the pupil in an aberration-free and vignetting-less system. (**B**), Optical invariant for the collection of emission light (*I*_c_). The FASHIO-2PM has a twofold larger angle of the cone of light at the image plane than that of excitation light. To maximally utilize the large *I*_c_, *r_s_* and/or *θ*_s_ should be large.

To achieve fast imaging, we decided to use a resonant-galvo scanning system instead of a galvo-galvo system. To maximize *r_m_* sin*θ*_m_ in Eq. (1), we chose a resonant mirror with a 7.2 x 5.0 mm elliptical clear aperture, 26-degree maximum angle and 8 kHz resonant frequency (CRS8KHz, Cambridge Technology). These settings achieve a 2.5 times higher optical invariant than that obtained with a 10-degree angle and at 12 kHz (CRS12KHz)(*6*) and twofold faster imaging than that achieved under 12 x 9.25 mm, 26 degrees, and 4 kHz (CRS4KHz)(*5*).

We sought to develop a new objective lens with a large pupil capable of supporting an optical invariant higher than the optical invariant of CRS8KHz. We further aimed to make the pupil larger to collect as much fluorescent emission light as possible from a specimen. The optical invariant for the collected light *I*_c_ of the LS2PM is determined independently from *I_e_* as follows:

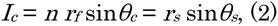

where *θ_c_* is the angle of the cone of the collected light from the neurons and *r_s_* and *θ_s_* are the sensor radius and angle of the cone of light exiting the collection optics, respectively (Fig. 1B). The angle *θ_c_* is designed to be twice as large as *θ_e_* in our system(*6*) This larger angle of *θ_c_* leads to *I_c_* > *I_e_* and can improve the SNR. To maximally utilize the large *I*_c_, *r_s_* and/or *θ_s_* should be large. A condensing lens can be utilized to increase *θ*_s_ by placing it before a conventional GaAsP PMT that has a small *r*_p_(*6*). We did not, however, follow this strategy because the output current of the PMT, which positively correlates with the SNR, is low. Instead, we developed a large-aperture *r_s_*, high output current, and high-sensitivity GaAsP PMT.

### Detailed design

In a laser scanning microscope, the laser radius *r_m_* is restricted by the utilizable area of the scanning mirror. We therefore selected a resonant mirror with a high precision flatness of < 0.015λ rms at 633 nm across the entire mirror surface. Second, we expanded the laser radius *r*_m_ to 2.75 mm (1/e^2^) using a beam expander and finally projected the expanded laser onto the selected high-precision resonant mirror. (Note the tradeoff between the increase in the mirror area on which the laser beam is projected and the wavefront error. Because the flatness on the outer periphery of the mirror is, in general, lower than that in the center, such usage of a large reflection area of the mirror considerably increases the amount of wavefront error, i.e., the deviation between the ideal wavefront and the system wavefront, leading to decreases in the efficiency of 2P excitation and in the optical resolution). We increased the scan angle of the mirror *θ*_m_ from ∼15 degrees (Nikon A1R MP) to 25.3 degrees at a resonant frequency of 8 kHz. Thus, the *I*_e_ of our system is 0.60, which is significantly higher than that of conventional 2P microscopes (*Ie of the Olympus FVMPE-RS with the XLPLN25XWMP2 objective lens (1.05 NA and ∼ 0.5 x 0.5 mm2 FOV) is ∼ 0.267, see also Bumstead 2017 for other microscopes*).

To ensure that the optical invariant at the objective is equal to or greater than 0.6, we designed a large objective lens that does not fit with a commercial-standard microscope (e.g., Leica, Nikon, Olympus and Zeiss) (Fig. 2A, see also Table S2 for lens specifications). The NA for excitation of the objective was determined to be 0.4 to achieve a single-cell resolution along the z-axis(*15*). Because *I*_e_ = *n r*_f_ sin*θ*_e_, the FOV is 3 x 3 mm^2^, which is sufficiently large to monitor multiple brain regions. The objective lens had a 56 mm pupil diameter (dry immersion; 4.5 mm working distance, 35 mm focal length). The tube and scan lenses were also designed to satisfy the optical invariant (0.6) (Fig. 2A, see also Table S2 for lens specifications). The 2.75 mm radius laser *r_m_* was projected onto the entrance pupil of the objective through a relay system including the scan and tube lenses, resulting in an *r*_p_ of 14 mm in radius. For collection, we further increased the angle *θ_c_* to collect as much fluorescence from the sample as possible, resulting in an NA of 0.8. Finally, our objective had a rear aperture diameter of 64 mm, a 170 mm height with an 84 mm diameter (without a flange) and a weight of 4.2 kg. The optical invariant of the objective for collection was 1.2.

**Fig. 2.**
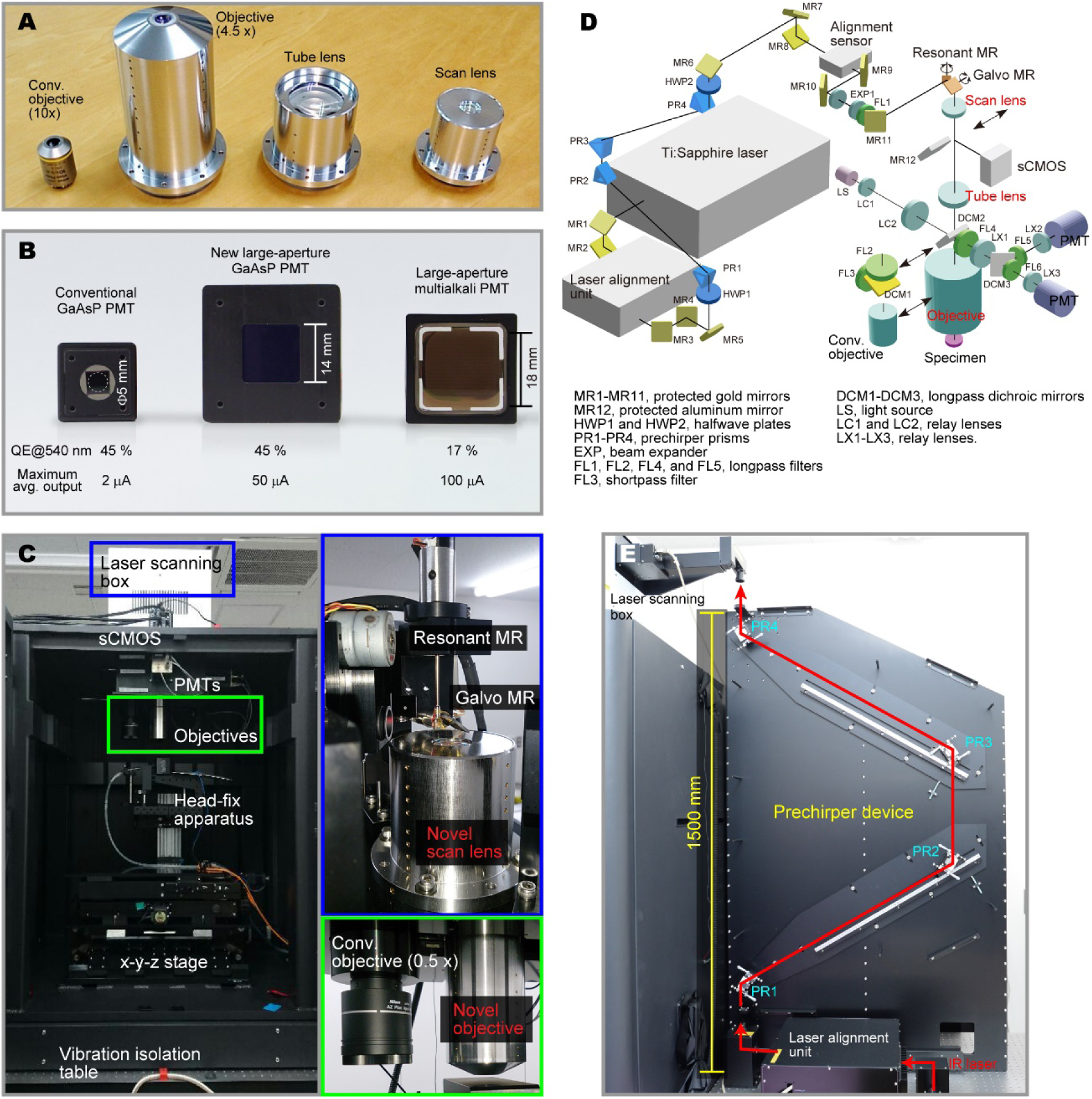
Design and optical layout of the FASHIO-2PM. (**A**) Conventional objective lens (CFI Plan Fluor 10X/0.30, Nikon) and newly developed large objective, tube and scan lenses. (**B**) Conventional GaAsP PMT (left), newly developed large-aperture GaAsP PMT (middle), and conventional large-aperture multialkali PMT (right). (**C**) Overall image of the microscope (left, front view), inner structure of the laser scanning box (blue square, top right), and enlarged image of the objective lenses (green square, bottom right). (**D**) Optical layout of the FASHIO-2PM (see Results and Methods for details). A set of conventional objectives, i.e., DCM1, FL2 and 3, and MR12, are manually switchable (arrows) for one-photon imaging with LS and sCMOS (see Methods). (**E**) Prechirper device (side view of the microscope) including 4 prisms (light blue) with a single path to realize alignment stability. It compensates for the IR laser pulse width (red arrow) broadening that occurs in optics (the maximum amount of compensation for group delay dispersion = ∼14400 fs^2^ at 900 nm).

To maximally utilize the optical invariant, we developed or used a large-aperture (*r*_s_) high-sensitive photodetector, a new GaAsP PMT (14 mm^2^ aperture, see also Fig. S1 for specification) or a commercially available large-aperture multialkali PMT (18 mm^2^) (Fig. 2b;). The new GaAsP PMT has a large aperture, ∼10x larger than that of conventional GaAsP PMTs (*Φ* 5 mm), and a QE that is ∼2.6x higher than that of the multialkali PMT (45% vs. 17% at 550 nm). Another advantage of the new PMT is a significantly higher maximum average output current than the conventional current (50 μA vs. 2 μA). The large current will contribute to an increase in the SNR and in the upper limit of the signal dynamic range such that one can monitor neurons with low-and high-fluorescence signals at the same time during fast imaging.

For the system (Fig. 2C and D), an infrared (IR) laser from a laser generator is introduced into a laser alignment unit and a prechirper device consisting of four prisms (Fig. 2E) to avoid the degradation of pulse stretching of the laser pulses(*16*) due to the group delay dispersion of the optics. The IR laser is then led to the resonant-galvo scanning system and the pupil of the objective lens through the scanner, tube lenses, and a dichroic mirror. The IR laser finally illuminates a specimen. The emission light from it is collected by the objective and is led to PMTs via dichroic mirrors. The power transmission ratio through the entire system is ∼25%. Importantly, the scanning angle of a resonant mirror can be increased while maintaining a constant resonant frequency, permitting fast imaging. Conversely, as another tradeoff, it decreases the pixel dwell time (the time that the laser dwells on each pixel position: FASHIO-2PM, ∼18-36 ns; Nikon A1R MP, ∼70-140 ns), which decreases the SNR. Thus, to overcome this tradeoff, we used the new large-aperture GaAsP PMT with higher quantum efficiency than that of the multialkari PMT. For the excitation system, a simulation of the encircled energy function (EEF), representing the concentration of energy in the optical plane, showed that 80% of the energy of light in the entire FOV is contained within a radius of 1.1 μm (Fig. 3A and B; see also Fig. S2 for other parameters). This value, even at the edge of the FOV, is almost equivalent to the diffraction limit, indicating high efficiency of the two-photon excitation and a high spatial resolution on all three axes across the entire FOV. The difference in excitation energy between the center and edge of the FOV was designed to be less than 1%, thereby preserving uniform excitation within the entire FOV. Our objective lens has a superior SR, which is the index of the quality of the point spread function (PSF), of ∼0.99 over the FOV, indicating a practically aberration-free objective. The image resolution was estimated based on the lateral and axial full widths at half maximum (FWHMs) of the bead images (Fig. 3C). The lateral FWHM of the bead images was 1.62 ± 0.07 μm (s.e., n = 14) and 1.62 ± 0.04 μm (n = 17), and the axial FWHM was 7.59 ± 0.19 μm and 10.22 ± 0.14 μm with compensation (see Methods) at ≤ 100 μm and at 500 μm, respectively, below the surface of the cover glass, indicating single-cell (soma) resolution on the x-, y-and z-axes.

**Fig. 3.**
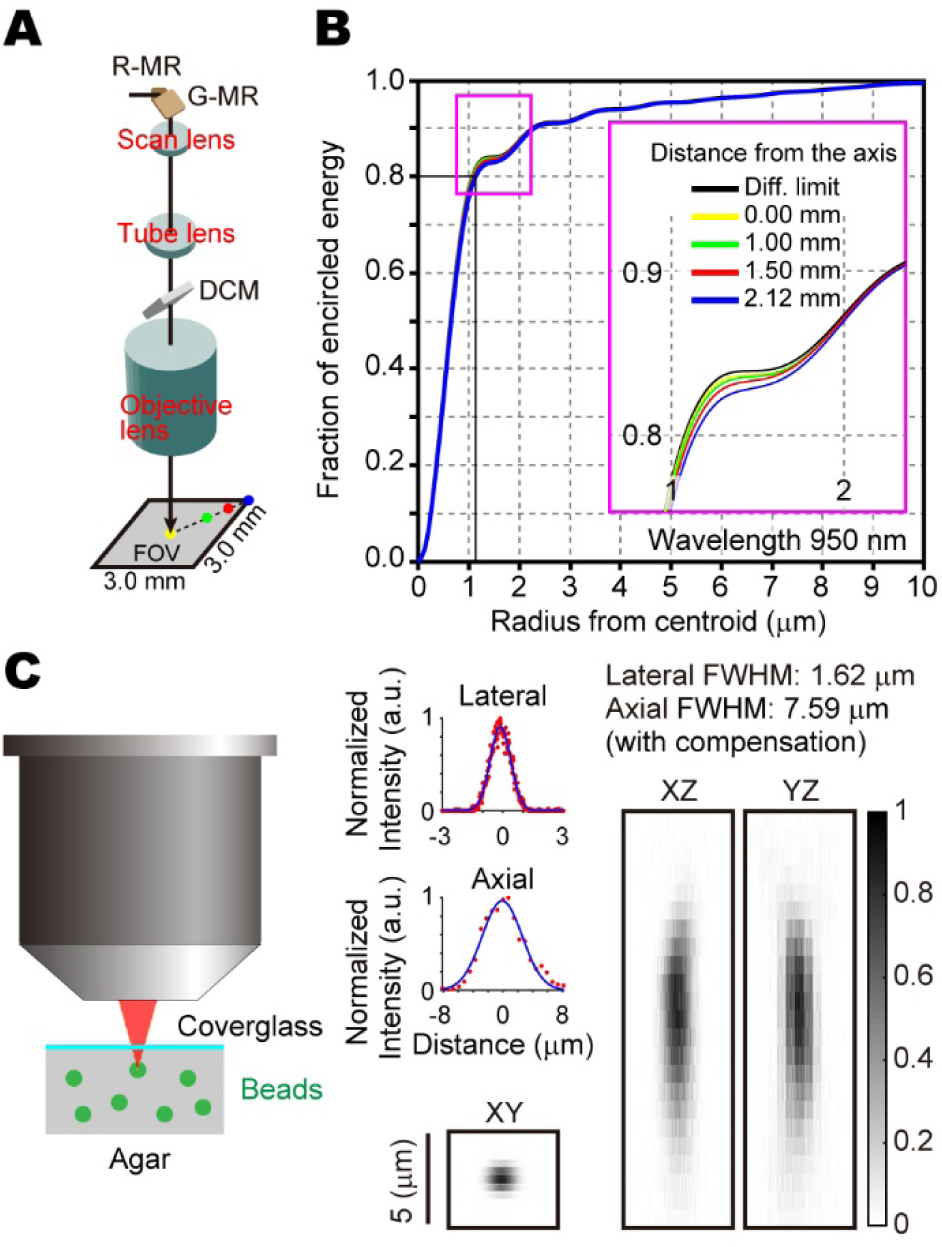
Optical performance of the FASHIO-2PM. (**A, B**) Simulation of the encircled energy function (EEF). (**A**) The EEF was simulated for a system including resonant and galvo mirrors, scan and tube lenses, a dichroic mirror and an objective. The actual distances between these components were used for the simulation. (**B**) The result of the simulation. The black line indicates the diffraction limit. Colored lines indicate the results from the axis (yellow) to the edge of the FOV (blue). Inset, expanded area (magenta in **A**) (see also Fig. S2 for other parameters). Our objective lens has SR of ∼ 0.99 over the FOV. (**C**) The point spread function (PSF) profile of the system, measured using fluorescent beads. Radial and axial excitation PSF measurements were performed at ≤ 100 μm below the surface of the cover glass. FWHM, full width at half maximum of the bead images (0.5 and 1.0 μm in diameter).

### In vivo Ca^2+^ imaging: proof-of-concept experiments

To demonstrate the quality and scale of FASHIO-2PM imaging compared to imaging with a conventional two-photon microscopic system, we expressed the genetically encoded calcium indicators (GECIs) G-CaMP7.09(*17*) or GCaMP6f(*18*) in neurons widely distributed across the cortical area and performed wide- and contiguous-FOV two-photon Ca^2+^ imaging. Specifically, we injected an adeno-associated virus (AAV)-conjugated Ca^2+^ indicator (AAV-DJ-Syn-G-CaMP7.09 or AAV9-Syn-GCaMP6f) into the cortex or lateral ventricle of a wild-type mouse on postnatal day 0-1 (P0-1) (Fig. 4A; see also Methods), opened a large cranial window (∼4.5 mm in diameter), and monitored the Ca^2+^ activity after P28. To estimate the number of neurons labeled with this injection, we stained cortical slices including the primary somatosensory cortices of the forelimb (S1FL) and hindlimb (S1HL) regions, the primary motor cortex (M1), the posterior parietal cortex (PPC), and the barrel cortex with antibodies against NeuN, a neuronal marker, and GAD67, an inhibitory neuron marker. We found that 85.1-90.2% of all layer 2/3 (L2/3) excitatory neurons (i.e., non-GABAergic neurons) in these cortical areas were labeled through this injection (Fig. 4A-C; see also Methods for the cell estimation procedure).

**Fig. 4.**
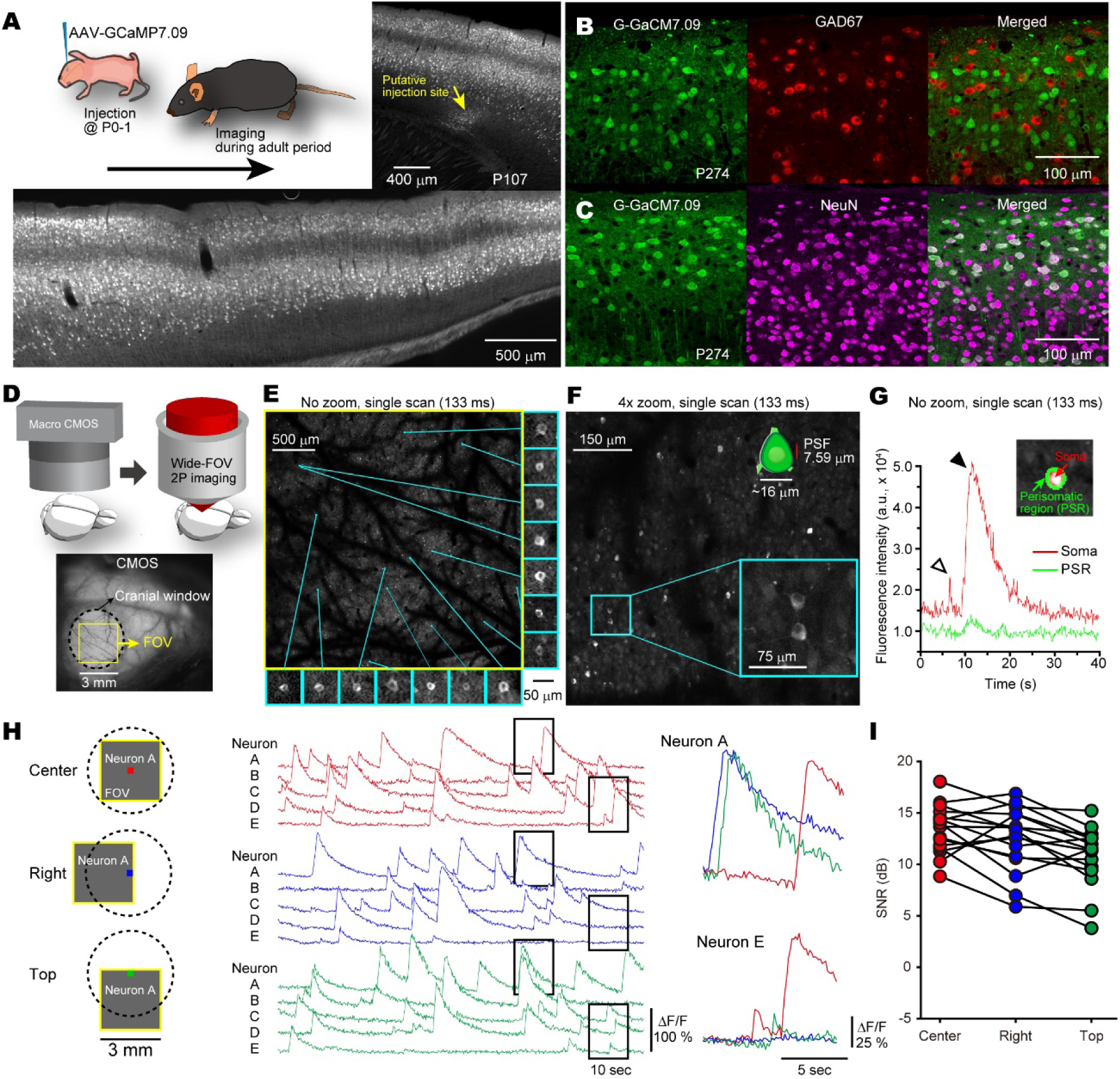
In vivo Ca^2+^ imaging from cortical excitatory neurons in layers 2/3. (**A**) Schematic illustration of the experimental protocol for the injection of an adeno-associated virus (AAV)-conjugated Ca^2+^ indicator (G-CaMP7.09 or GCaMP6f). Sagittal brain slices from an adult mouse were monitored to clarify the labeled cortical neurons and the putative injection site. (**B**) Representative cortical slice with G-CaMP7.09 (green) and GAD67 as a marker for GABAergic neurons (red). (**C**) Representative cortical slice with G-CaMP7.09 (green) and NeuN (magenta). (**D**) Schematic illustration of the experimental approach for Ca^2+^ imaging. Macroscopic one-photon CMOS imaging showing a craniotomy (dashed line) and the FOV (yellow) for two-photon imaging on the right hemisphere. (**E**) Ca^2+^ imaging of a contiguous full FOV (3 × 3 mm^2^) including layer-2 cortical neurons labeled with GCaMP6f at 7.5 Hz sampling rate. One frame image without averaging was shown. Representative neurons are shown in the blue boxes. (**F**) Ca^2+^ imaging with a 4× magnification to clarify the nuclei of the neurons (blue box). One frame image without averaging was shown. The inset presents a size comparison between a soma and the PSF. (**G**) Representative Ca^2+^ transients at the soma (red) and fluorescence changes in the perisomatic region (green). The inset shows the manually selected region of interest (ROI). (**H**) Left: Schematic diagram of the assessment of the signal-to-noise ratios (SNRs) as recorded from the center (red), right (blue) and top (green) of the FOV. Middle: Representative Ca^2+^ signals from neurons A-E. Right: Magnified signals of neurons A and E (black boxes in the middle plots). (**I**) Summary of the SNRs recorded from the three regions of the FOV (n = 16 neurons).

The full 3 ⨯ 3 mm^2^ FOV (2048 x 2048 pixels), including the somatosensory area, was scanned at 7.5 frames/s (G-CaMP7.09; see Video S1 for raw and ΔF/F data representations). To image L2/3 neurons, we used ∼60∼80 mW of laser power with the GaAsP PMT (< 180 mW with the multialkali PMT) at the front of the objective. This power level is below 250 mW, which may initiate heating damage or phototoxicity in conventional-FOV microscopes(*19*).

Because the PSF on the z-axis was 7.59 μm, almost equivalent to half the diameter of L2/3 neurons, we were able to detect the nuclei as intracellular structures devoid of GCaMP6f fluorescence within the cell bodies, even at the edge of the FOV (Fig. 4D and E). The nuclei were also confirmed by the 4⨯ zoom imaging mode, in which the optical x-y-z resolution was kept the same as in the non-zoom mode but the pixel x-y resolution was increased (FOV: 0.75 ⨯ 0.75 mm^2^; x-y pixel size: 0.366 μm) (Fig. 4F). These results demonstrate a sufficient spatial resolution for single cells along all axes throughout the entire FOV, as supported by the EEF simulation (Fig. 3A and B). We manually selected regions of interest (ROIs) on the somata and perisomatic regions (PSRs) of single neurons and were able to monitor larger Ca^2+^ transients with both fast (open arrowhead in Fig. 4G) and long (filled arrowhead) decay times at the soma compared to those in the PSR. Importantly, because of the very low field curvature and F-theta distortion (< 4 μm and < 1 μm across the FOV, respectively; see Fig. S2), we were able to continue monitoring the same neurons and the same geometric structure composed of a large number of neurons while changing the FOV. We also compared the SNRs among Ca^2+^ signals monitored from the same neurons at the center, right and top of the FOV (Fig. 4H). Although the SNR at the edges of the FOV (i.e., the top and right) was slightly lower than that at the center (Fig. 4I), it was still sufficient to identify Ca^2+^ transients. This finding demonstrates that the FASHIO-2PM is able to monitor Ca^2+^ activity with a high SNR throughout the full FOV. We were also able to monitor Ca^2+^ activity from deep-layer neurons (500 μm below the cortical surface) with a sufficient SNR (Fig. S3 and Video S2).

### Functional network analysis with single-cell resolution

Wide and contiguous scanning of FASHIO-2PM enables estimation of functional connectivity among a large number of neurons, leading to reliable investigation of functional network architecture. We monitored Ca^2+^ activity of ∼16000 cortical neurons from layer 2 (100–200 μm below the cortical surface) spanning 15 sensory-motor and higher-order brain areas during head-fixed awake mice. (Fig. 5A). An algorithm called low-computational-cost cell detection (LCCD)(*20*) was applied to extract the Ca^2+^ activity from each neuron (Fig. 5B for clarification of ROIs and neurons, see also Methods for the section “Image analysis”, Fig. 5C for examples of Ca^2+^ activity randomly selected from the ROIs). We were able to monitor movement-related and unrelated spontaneous Ca^2+^ signals from a large sample (Fig. 5D).

**Fig. 5.**
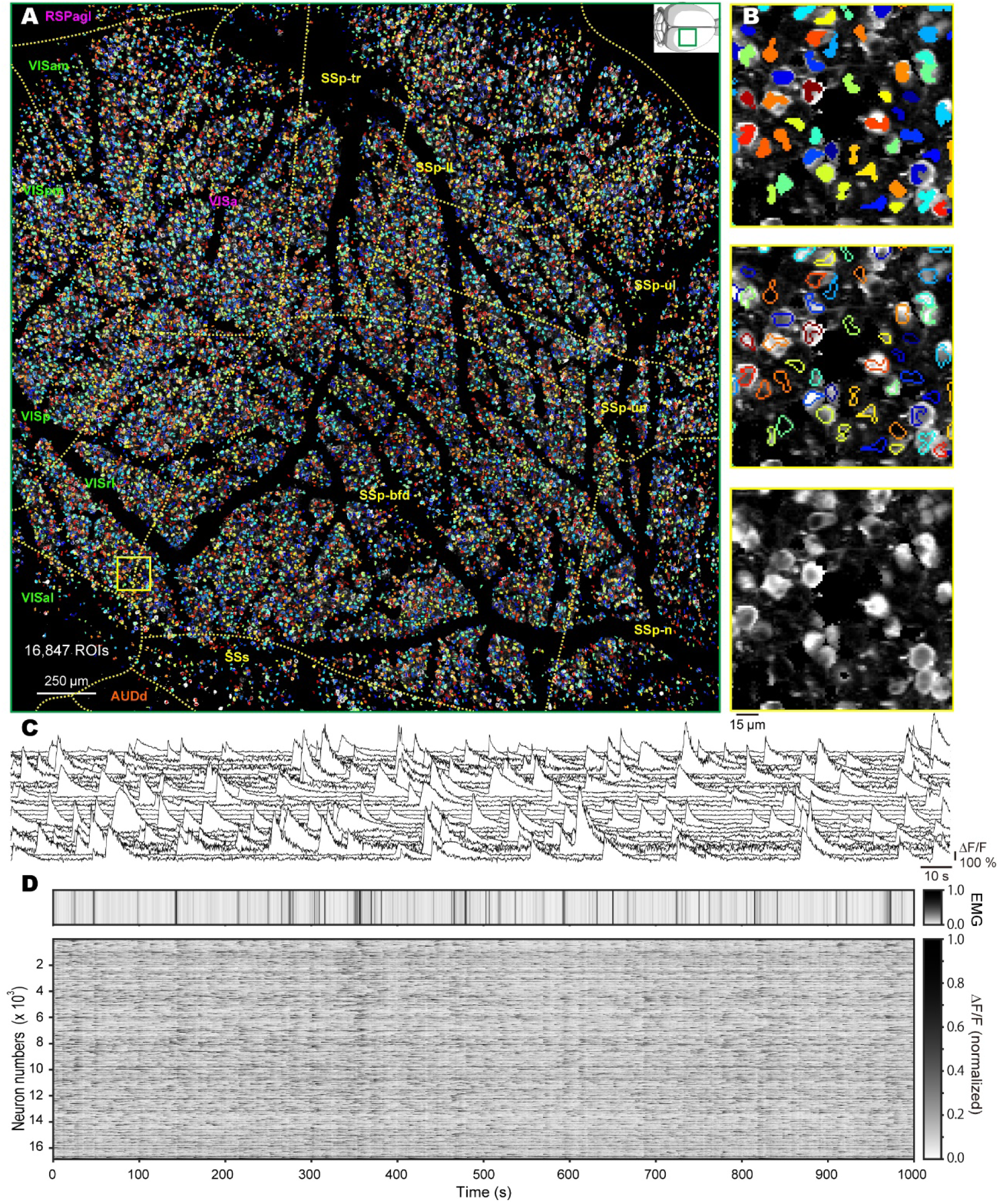
Automated cell detection with the low-computational-cost cell detection algorithm and neural activities from 15 areas. (**A**) ROIs detected via the low-computational-cost cell detection (LCCD) algorithm (120 μm depth from the cortical surface; 3 × 3 mm^2^ FOV; 2,048 x 2,048 pixels; 7.5 Hz sampling rate). The total number of ROIs is 16,847. Individual ROIs are randomly colored for visual clarity. Inset, a mouse brain and the FOV shown as a green box. (**B**) Magnified view of the area shown as a yellow box in (**A**), with filled ROIs (top), with open ROIs (middle) and without ROIs (bottom), to clarify the shapes of the ROIs and neurons. (**C**) Ca^2+^ signals from randomly selected 25 neurons in the area shown on the right in (b) for clarification of the signals. (**D**) Electromyography (EMG) normalized between 0 to 1, and Ca^2+^ signals normalized between 0 to 1 in each neuron. Cells are sorted in descending order of decoding accuracy. Overlaid yellow lines show borders according to the Allen Mouse Common Coordinate Framework. Briefly, SSp-n, S1-nose; SSp-bfd, S1-barrel field; SSp-ll, S1-lower limb; SSp-ul, S1-upper limb; SSp-tr, S1-trunk; SSp-un, S1-unassigned area; SSs, S2; VISal, anterolateral visual area; VISam, anteromedial visual area; VISp, V1; VISpm, posteromedial visual area; RSPagl, retrosplenial area, lateral agranular part; VISa, anterior area; VISrl, rostrolateral visual area.

We measured pairwise partial correlation coefficients (PCC) between the Ca^2+^ activity with high SNR (see Methods for ROI selection), which removes false associations in Ca^2+^ activities that were derived from animal movements (Fig. 6A and Methods for the partial correlations). We then examined the distribution of PCCs corresponding to physical distance between neurons (Fig. 6B) and found long-range pairs that cannot be observed in a small FOV of a conventional microscope (Fig. 6C).

**Fig. 6.**
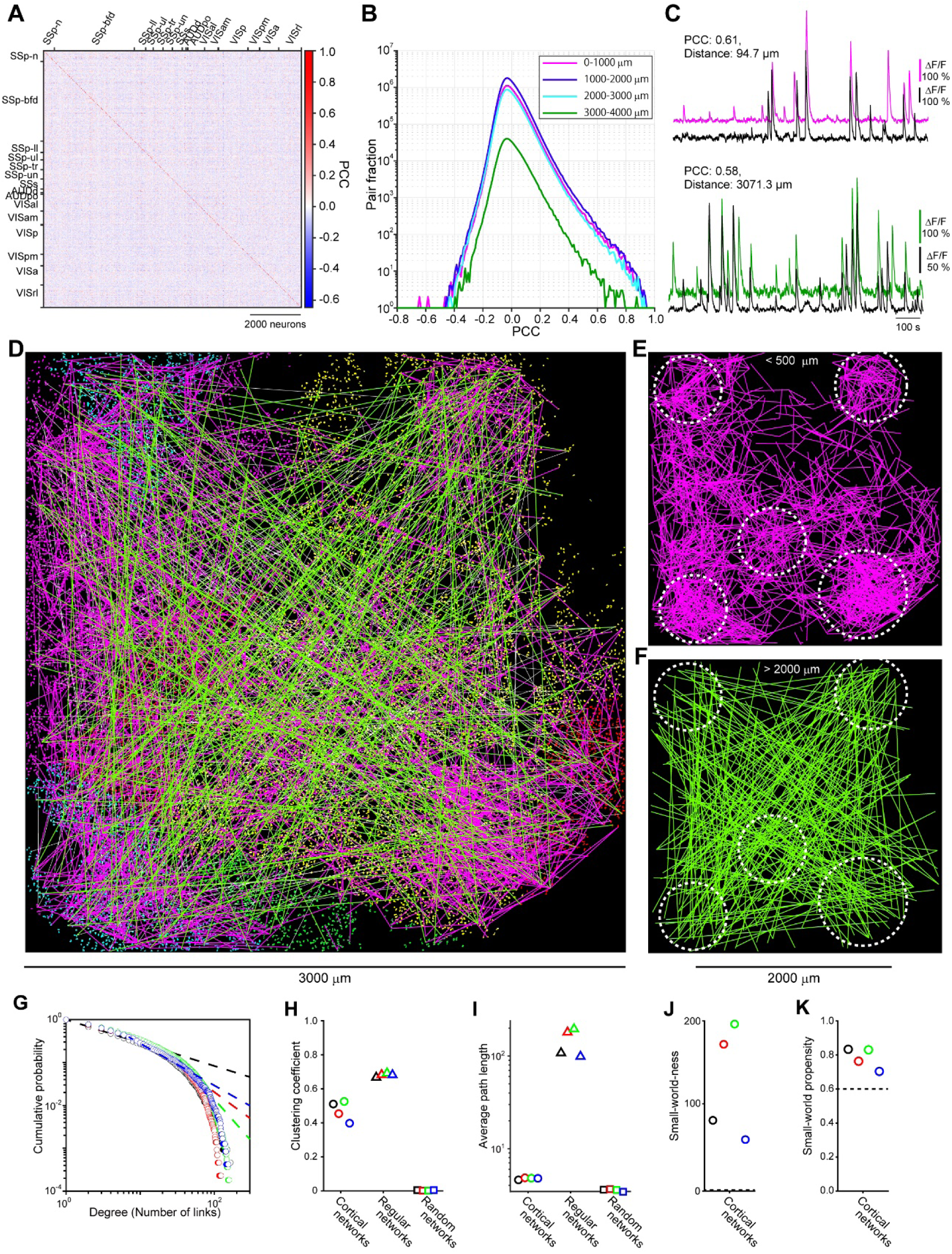
Correlation-based functional network analyses. (**A**) Partial correlation coefficient (PCC) matrix for a representative mouse. Correlations between the neural activities and the gyro sensor signal have been partialled out. (**B**) Histogram of PCC depending on pair’s distances. (**C**) Examples of pairs of Ca^2+^ signals that show short-and long-distance correlations. (**D**) Functional network on cortical map. 2000 links (∼5.6% of total links) were randomly selected for clarification of the network structure (1000 green links for > 2000 μm; 1000 magenta links for < 500 μm distance pair) and mapped on ROIs. Different ROI colors represent different cortical regions. (**E, F**) Extracted short-(**E**) and long-distance (**F**) links shown in (**D**) for clarification. The locations of cluster-like short-link populations are shown in the white circles. (**G**) Cumulative degree distributions of the cortical networks. The dashed lines represent the best-fit power laws estimated via maximum likelihood fitting with the Kolmogorov–Smirnov minimization approach(*21, 24*). (**H, I**) Clustering coefficients (**H**) and average shortest path lengths (**I**) of the observed cortical, regular, and random networks containing comparable numbers of nodes and links. (**J, K**) Small-world-ness(*25*) (**J**) and small-world propensity(*26*) (**K**)of the cortical networks. The dashed lines represent the thresholds for indicating a small-world topology (1 in **J** and 0.6 in **K**). In (**G-K**), different colors correspond to different mice.

We regarded individual neurons and connectivity with PCC above 0.4 between neurons as nodes and links, respectively, resulting in the construction of a functional and binary undirected network that were neither too sparse nor too dense for assessment of network architecture(*21*) (see Methods). By mapping the network depending on the pair’s distances on the cortical map, we found that the links spanned multiple cortical regions in the short and long distances (Fig. 6D). The short links are likely to form cluster-like populations (Fig. 6E, with white circles denoting the populations), and the long links are likely to bridge the populations (Fig. 6F).

To quantitatively assess these observations, we calculated following network measures; number of links per node (i.e., degrees), the ratios of the numbers of triangles among the linked nodes (i.e., clustering coefficients), and the average shortest path lengths over all pairs of nodes (i.e., average path lengths). These measurements are known to characterize two ubiquitous architectures in real-world networks: scale-free architecture which is characterized by the existence of a small number of high degree nodes(*22*) and small-world architecture which is characterized by a high clustering coefficient and a short average path length(*23*). To the best of our knowledge, verification of whether the brain has either the small-world or scale-free properties, both or neither has yet to be demonstrated using a large number of single-cell activities from multiple brain regions. To evaluate assess a scale-free property in the cortical networks, we mapped the degree distributions of the networks and examined whether the distributions exhibit power-law attenuation (Fig. 6G, dashed lines). A series of statistical tests proposed in previous studies(*21, 24*) showed that the cortical networks do not (or only weakly) satisfy scale-free architecture (Table S4, see also Methods). Nonetheless, we found very few hub-like neurons (0.1-0.01% of neurons used for analysis) that have ∼10^2^ or more links in the cortical network (Fig. 6G). To evaluate a small-world property in the cortical networks, we compared clustering coefficients and average path lengths with those of random and regular networks containing a comparable number of nodes and links. We found that the cortical networks showed significantly higher clustering coefficients than random networks (Fig. 6H and Table S5) and shorter path lengths than regular networks (Fig. 6G and Table S5). These results are evidence of small-world architecture(*23*), which is further supported by two small-world metrics proposed in previous studies; the small-world-ness(*25*) was significantly higher than the criterion of 1 (Fig. 6J), and the small-world propensity(*26*) was higher than the criterion of 0.6 (Fig. 6K).

## Discussions

In this study, we have described the FASHIO-2PM, which is based on the novel approach of combining a large objective lens with a resonant-galvo system for fast and wide-FOV two-photon microscopy and achieves a 36-fold increase in the overall imaging area compared to a conventional 2P microscope that is used in neuroscience (i.e., 9 mm^2^ vs. 0.25 mm^2^ FOV). We emphasize that our microscope concurrently achieves all of the following key benchmarks: 1) contiguous scanning, 2) a wide FOV, 3) single-cell resolution, 4) a fast sampling rate, 5) practically aberration-free imaging and 6) a high SNR.

One of the fundamental differences in two-photon image acquisition between our microscope and others is whether the fast sampling speed applies over a wide FOV while suppressing aberrations. This achievement of the FASHIO-2PM is primarily based on the performance of the newly designed objective. Regardless how fast the imaging speed is, if the SNR is inadequate, the speed must be decreased until the SNR is sufficient. Thus, increasing the performance of the objective lens is critical for achieving fast imaging with a sufficient SNR. Other mesoscopes (including the two-photon random access mesoscope, i.e., 2P-RAM, and the Trepan2p) use a higher NA for excitation (0.6 and 0.43, respectively) than that used by ours (0.4). However, not every high-NA objective lens provides higher-resolution and brighter images than those achieved by a lens with a lower NA. The important point is how efficiently the laser can excite fluorescent proteins (e.g., a Ca^2+^ sensor). In other words, the number of photons must be increased as much as possible within a focus area that is as small as possible by suppressing aberrations. Our system exhibits a superior EEF that is close to the diffraction limit, indicating highly efficient excitation. Although we cannot completely compare the systems, the simulation results were supported by the fact that the PSF value we measured by the FASHIO-2PM with a 0.4 NA shows significantly higher resolution on the z-axis than that obtained by the Trepan-2P with a 0.43 NA (7.59 μm at ≤ 100 μm vs. ∼12 μm at 55 μm; 10.22 μm at 500 μm vs. ∼12 μm at 550 μm from the imaging surface, respectively). The SR of our objective (∼0.99 across the FOV) indicates more efficient excitation than that of another large objective (> 0.8 in 2P-RAM(*6*)). Thus, even with 1/3 to 1/4 the pixel dwell time of other mesoscopes, our microscope achieves fast imaging with sufficient SNRs for the entire FOV. Another critical factor affecting the capability of fast imaging is the NA for collection (0.8 NA in the FASHIO-2PM vs. 0.43 NA in the Trepan-2P). Overall, our system with the aberration-free large objective lens can realize fast, wide-FOV imaging with a sufficient SNR.

Our microscope also differs from other 2P microscopes when monitoring a large number of neurons. For this purpose, one uses an image tiling technique involving the subdivision of a large x-y plane(*27*), the separation of a 3D volume into multiple smaller areas(*28–30*) or noncontiguous multiarea imaging(*5–7*) (e.g., 0.25 mm^2^ x 4 areas). Of course, each microscopy approach has certain advantages and disadvantages. For example, the current version of the FASHIO-2PM cannot achieve fast volumetric imaging, which is useful for investigating, for example, the operation mechanisms of a single cortical column (∼1 mm^2^) by monitoring multiple layers(*28*–*30*). Instead, our microscope can monitor cortical-wide interactions from a contiguous FOV (9 mm^2^) that contains ∼15 brain regions.

This wide and contiguous imaging offers the opportunity to measure the correlations of Ca^2+^ signals between thousands of neurons (Fig. 6). We here constructed correlation-based functional networks at a single-cell resolution and showed that the cortical networks exhibit only weak scale-free but a significant small-world architecture. Our large-scale observation makes statistical analyses of network architecture (*21, 24*) applicable, and thus providing more reliable assessments than previous investigations with a smaller number of samples (ex. 24 neurons(*31*)). By expanding a conventional FOV, we discovered long-distance correlations between neural activities that have been thought to be quite rare cases or pairs from background small noise but not neurons when conventional small FOV imaging is used(*32*). These correlations, potentially contribute to the small-world architecture at single-cell level. Further studies are needed to elucidate how the network emerges, and relates to brain functions and animal behaviors. Monitoring a large network with its elements including hub-like rare neurons that we found will enable revealing biological dynamics in detail.

In the future, our microscope can be improved by incorporating novel features that have been developed for other microscopes. Because it is designed to have a simplified optical path, the FASHIO-2PM offers the potential to install a photostimulator for the manipulation of neural activity(*33*) as well as components for 3D(*28–30*), multicolor(*34*), and three-photon imaging(*35*). Notably, Han and colleagues(*30*) developed a two-color volumetric imaging system to monitor the neural activity of cortical columns. Because this technique is compatible with other imaging systems, it can be implemented in our microscope; thus, L2/3 and L5 neurons will be monitored at the same time with different colors. In addition to these improvements on the hardware side, additional software techniques can also be implemented in the FASHIO-2PM, such as cell-type-specific(*36, 37*) or subcellular-component-specific(*38*) imaging with various combinations of transgenic mouse lines(*39*) and GECIs(*18, 40*), fast Ca^2+^ sensors to follow single action potentials(*34*), or an algorithm for spike detection from Ca^2+^ signals(*41, 42*). Moreover, various electrical signals(*4, 43–45*) can be simultaneously recorded to produce synergistic effects or provide complementary information to resolve questions about cortical dynamics and enable the discovery of new phenomena underlying brain functions.

## Material and method

### FASHIO-2PM

For two-photon Ca^2+^ imaging with G-CaMP/GCaMP, a Ti:sapphire laser (Mai Tai eHP DeepSee, Spectra-Physics) was tuned to 920 nm. Resonant and galvanometric mirrors were used for laser scanning (CRS-8kHz and VM500, respectively, Cambridge Technology). Emission light was collected through 500 nm and 560 nm long-pass dichroic mirrors (DCM2 and 3 respectively, shown in Fig. 1) and 515 - 565 nm (FL5) and 600 – 681 nm (FL6) band-pass emission filters with Multialkali photomultiplier tubes (PMTs) (R7600U-20, Hamamatsu Photonics KK) or GaAsP PMTs (Hamamatsu Photonics KK). The PMT signals were pre-amplified and digitized using an analog-to-digital converter connected to a PC. The images (3 × 3 mm) were recorded at 7.5 Hz using a resolution of 2,048 × 2,048 pixels, with a software program (Faclon, Nikon Instruments Inc.). For one-photon macroscopic imaging, we used a blue LED light source (center wavelength: 465 nm, LEX2-B-S, BrainVision Inc.) coupled with an optical bundle fiber and a 500 nm short-pass excitation filter for illumination. A 500 nm long-pass dichroic mirror reflected the excitation light. The light illuminated the skull and blood vessels on the cortex through a conventional objective (AZ Plan Apo 0.5×, NA: 0.05/WD: 54 mm, Nikon). Emission light was collected through the dichroic mirror and a 525 nm long-pass emission filter with a sCMOS camera (Zyla 5.5, Andor).

### Animals

All animal experiments were performed in accordance with institutional guidelines and were approved by the Animal Experiment Committee at RIKEN. Wild-type mice (C57BL/6JJmsSlc, Japan SLC, Shizuoka, Japan), CAG-lox-stop-lox-tdTomato mice (Jackson Labs stock #007905) crossed with VGAT-cre mice(*46*) and Rbp4-Cre (MMRRC stock #031125-UCD) were used. Both male and female mice were used indiscriminately throughout the study. In all experiments, the mice were housed in a 12 h-light/12 h-dark light cycle environment with ad libitum access to food and water.

### Adeno-associated virus (AAV) vector preparation

G-CaMP7.09(*17*) was subcloned into the synapsin I (SynI)-expressing vector from a pN1-G-CaMP7.09 vector construct. pGP-AAV-syn-jGCaMP7f-WPRE(*47*)) was F from the Douglas Kim & GENIE Project (Addgene plasmid # 104488; http://n2t.net/addgene:104488; RRID:Addgene_104488). The following adeno-associated viruses (AAVs) were produced as described previously(*48*): AAV-DJ-Syn-G-CaMp7.09. AAV9-Syn-GCaMP6f (#AV-9-PV2822) and AAV9-Syn-Flex-GCaMP6s (#AV-9-PV2821) were obtained from Penn Vector Core. These AAVs were also donated from the Douglas Kim & GENIE Project.

### Virus injection

AAVs were injected into the neonatal lateral ventricle(*49, 50*) or the neonatal cortex(*51, 52*). For the neonatal intraventricular injection, AAV9.Syn.GCaMP6f (Penn Vector Core; titer 7.648 × 10^13^ GC/ml) was used (Fig. 4). For the neonatal cortical injection, AAV DJ-Syn-G-CaMP7.09-WPRE (titer 2.8 × 10^13^ vg/ml) was used after being diluted with saline to 0.7–1.0 x 10^13^ vg/ml (Fig. 4, 5 and 6). All of the AAV solutions were colored with 0.1% Fast Green FCF (15939-54, Nacalai Tesque Inc.), where they then were filled in a glass pipette (7087 07, BRAND or Q100-30-15, Sutter Instrument) with a tip diameter of 50 mm and a tip angle that was beveledto 45° by the grinder (EG-401, NARISHIGE). P1 or P2 mice were collected from the cage and cryoanesthetized for 2-3 min before injection, then mounted in a neonatal mouse head holder (custom-made, NARISHIGE). The tip of the glass pipette was guided to the injection site using a micromanipulator (NMN-25 or SM-15R, NARISHIGE). The coordinates of the intraventricular injection site were approximately anteroposterior 1.5 mm, mediolateral 0.80 mm and dorsoventral 1.5 mm from lambda and scalp. The cortical injection site was located at a depth of 0.3–0.5 mm in the frontal area. The total volume of injected AAV solution was 2 and 4 μl for the intraventricular and cortical injection, respectively. After cryoanesthesia, the injection procedure was finished within 10 min and the pups were warmed until their body temperature and skin color returned to normal. Once the pups began to move, they were returned to their mother.

### Transcranial imaging

Transcranial imaging was performed 28–35 days after the AAV injection to examine the intensity and region of GCaMP expression. The mice were anesthetized with 2% isoflurane. Following anesthetization, the head hair of the mice was shaved and they were placed in head holders (SG-4N, NARISHIGE). Isoflurane concentrations during the procedure were maintained at 1–2%, and body temperature of the mice was maintained at 36–37°C with a heating pad (BWT-100, Bio Research Center). The scalp was incised along the midline, the skull was swabbed with cotton swab soaked with 70% (vol/vol) ethanol solution, and blood on the skull was washed with saline. A cover-glass (22 mm in diameter and 0.13-0.17 mm thick, Matsunami Glass Ind.) was placed on the skull. The brain was illuminated by a blue LED light with a center wavelength of 465 nm (LEX2-B, Brainvision) through a neutral density (ND) filter (NE06B or NE13B, Thorlabs) and a 506 nm dichroic mirror. Green fluorescence was corrected through a 536/40-nm filter. Fluorescence was recorded by a CMOS camera (MiCAM ULTIMA, Brainvision) using a software program (UL-Acq, Brainvision) under a tandem lens, objective lens (Ref. 10450030 Planapo 2.0×, Leica), projection lens (Ref. 10450029 Planapo 1.6×, Leica), and fluorescence microscope (THT-microscope, Brainvision). The field of view (FOV) was 8 × 8 mm^2^ (100 × 100 pixels). Every pixel collected light from a cortical region of 80 × 80 μm^2^. The exposure time was 10 ms. GCaMP expression was evaluated offline using various software programs (BV Ana, Brainvision and MATLAB, MathWorks). After transcranial imaging, the incised scalp was sutured with a silk suture (ER2004SB45, Alfresa Pharma Corporation).

### Surgery

The mice (13-26 weeks) were anesthetized with isoflurane (2%) for induction. Once the mice failed to react to stimulation, we administered hypodermic injections of the combination agent(*53*) – Medetomidine/Midazolam/Butorphanol (MMB) – at a dose of 5 ml/kg body weight for the operation. We dispensed MMB solution with medetomidine (0.12 mg/kg body weight), midazolam (0.32 mg/kg body weight), butorphanol (0.4 mg/kg body weight), and saline. We substituted fresh MMB solution every 8 weeks. After anesthesia with MMB, the head hair of the mice was shaved, and they were placed in head holders (SG-4N, NARISHIGE). During surgery, the body temperature of the mice was maintained at 36–37°C using a feedback-controlled heat pad (BWT-100, Bio Research Center), and their eyes were coated by ointment (Neo-Medrol EE Ointment, Pfizer INC.).

Our implant surgical operation fundamentally followed previously reported protocols(*54, 55*). After removing the scalp, the skull was swabbed with a cotton swab soaked into 70% (vol/vol) ethanol solution and 10% povidone-iodine (Isojin-eki 10%, Meiji Seika Kaisha). A 4.5-mm diameter craniotomy was then performed over an area that included the primary somatosensory area of the right hemisphere. The craniotomy was covered with window glass consisting of two different-sized micro cover-glasses (4.5 mm in diameter with 0.17-0.25 mm in thickness and 6.0 mm in diameter with 0.13-0.17 mm in thickness, Matsunami Glass Ind.). Beforehand, the smaller cover-glass was adhered to the center of the larger cover-glass using a UV-curing resin (Norland Optical Adhesive 81, Norland Products INC.). The smaller cover-glass was placed as close to the brain as possible, and the edge of the larger cover-glass was sealed to the skull with dental cement (Super Bond, Sun Medical). A stainless-steel head plate (custom-made, ExPP Co., Ltd.) was cemented to the cerebellum, as parallel to the window glass as possible. The exposed skull was covered with dental cement. Flexible wire cables were implanted into the neck muscles for electromyography (EMG) recording (Fig. 5, Fig. S5 and S6). After the surgery, the mice were dispensed a medetomidine-reversing agent, atipamezole hydrochloride (ANTISEDAN, Zoetis Inc.) solution at a dose of 0.12 mg/kg body weight; next, the mice were left under a heating pad and monitored until recovery.

### In vivo two-photon imaging

The mice (19-30 weeks) were anesthetized with isoflurane (2%) and an adhering electrode (SKINTACT, Leonhard Lang GmbH) was attached to provide electrical stimulation to the skin. We placed vibrating gyro sensors (ENC-03RC/D, Murata Manufacturing Co., Ltd.) to the left hind limbs (HLs) of the mice (Fig. 6). The mice were then moved to an apparatus (custom-made, ExPP Co., Ltd.) which was firmly screwed to the head plate that was cemented to the skull. The mouse bodies were covered with an enclosure to reduce large body movements as much as possible. The cover glass was swabbed with a cotton swab soaked in acetone. Before two-photon imaging, the cranial window was positioned at the center of the FOV for the sCMOS camera. This was due to the center of the FOV in the one-photon system corresponding with the FOV of the two-photon system.

All two-photon imaging was conducted with the FASHIO-2PM. The mice were awake during the calcium imaging. Recovery from the anesthesia was judged from the presence of mouse body movements measured by EMG or a gyro sensor (described in detail later). Imaging started 1 h or more after anesthesia was finished. The calcium sensor was excited at 920 nm and the calcium imaging parameters were as follows: 3.0 x 3.0 mm^2^ FOV; 2,048 x 2,048 pixels; 7.5 frame/s or 0.75 x 0.75 mm^2^ FOV; 2,048 x 2,048 pixels; 7.5 frame/s. The imaging depth below the pia was 100∼165 and 500 μm. For imaging L2/3 neurons, we used ∼60∼80 mW of laser power with the GaAsP PMT (< 200 mW with the multialkali PMT) at the front of the objective lens. For imaging L5 neurons, 300 mW of laser power was used with GaAsP PMT. The window glass was positioned parallel to the focal plane of the objective lens using goniometer stages (GOH-60B50 or OSMS-60B60, SIGMA KOKI CO., LTD.), which were under the head-fixation apparatus. This procedure was necessary to observe the neurons in the same layer of the cerebral cortex. The degree of parallelism was judged from two-photon images of the brain’s surface (3.0 × 3.0 mm^2^ FOV). All images were acquired using custom-built software (Falcon, Nikon), and the images were saved as 16-bit monochrome tiff files.

Mouse body movements were measured by EMG of the neck muscles or by the gyro sensor that was fitted to the left HL. The EMG signal was amplified 2,000 times and filtered between 10–6,000 Hz (Multiclamp 700B, Molecular Devices). HL motion was measured using a gyro sensor, and the signal was amplified and filtered using a custom-made analog circuit. The end times of each imaging frame were fed from the controller and were computed by the FPGA based on synchronization signals obtained from the resonant scanner. A pulse stimulator (Master-9, AMPI) controlled the two-photon imaging initiation and data logging. The EMG signals, gyro sensor signals, and the end times of each imaging frame were digitized at 20 kHz (Digidata 1440A, Molecular Devices).

### Image analysis

Brain motion of the recorded imaging data (8,000 frames) was corrected using the ImageJ plugin “Image Stabilizer”. We automatically detected active neurons using methods described within our studies(*20*). The regions of interest (ROIs) were detected within each of the 500 frames using the following 6 steps. (1) Removal of slow temporal trends and shot noise at each pixel. (2) Projection of the maximum intensity of each pixel intensity along the temporal axis. (3) Application of a Mexican hat filter to emphasize the edge of the cell body. (4) Adjustment of the contrast with the contrast-limited adaptive histogram equalization(*56*). (5) Binarization of the image using the Otsu method(*57*). (6) Detection of the closed regions whose pixel sizes were roughly equal to those of a single soma. Note that the final step effectively worked for preventing contamination through the inclusion of several neurons in a single ROI. After we completed the 6 steps, we integrated ROIs detected within every 500 frames, ensuring that several ROIs were not merged into a single ROI. The merge algorithm consisted of the following: (1) If the area of the overlap region between two ROIs occupied more than 40% of the whole area of either single ROI, we selected the larger one and deleted the smaller one. (2) If the area of the overlap region between the two ROIs occupied less than 40% of the total area of both ROIs, the overlap region was deleted from both ROIs. The ROIs obtained by our method with the parameters manually adjusted matched well with the ROIs detected by a constrained nonnegative matrix factorization (CNMF)(*42, 58*) from randomly selected small image sections.

The fluorescence time course *F_ROI_* (t) of each ROI was calculated by averaging all pixels within the ROI, and the fluorescence signal of a cell body was estimated as F(t) = *F_ROI_* (*t*) − *r* × *F_neuropil_*(*t*), with r = 0.8. The neuropil signal was defined as the average fluorescence intensity between 3 and 9 pixels from the boundary of the ROI (excluding all ROIs). To remove noise superimposed on F, we utilized the denoise method included in CNMF(*42*). The denoised F for 20 sec immediately after the start of imaging was removed until the baseline of the denoised F was stabilized. The ΔF/F was calculated as (F(t) − *F_0_*(*t*))/*F_0_*(*t*), where *t* is time and *F_0_*(*t*) is the baseline fluorescence estimated as the 8% percentile value of the fluorescence distribution collected in a ±30 sec window around each sample timepoint(*59*). We removed the ROIs that meet any one of the following conditions for topological analysis of functional network. (1) ROIs with low signal-to-noise ratio. We computed the signal-to-noise ratio of each ROI’s F using the “snr” function in MATLAB, where the signal was the ΔF/F below 0.5 Hz and the noise was the ΔF/F above 0.5 Hz. The ROIs with a signal-to-noise ratio below 3.0 were removed for data analysis. (2) ROIs on blood vessels. All analyses to detect blood vessels were performed in ImageJ. A median image of two-photon images for 500 frames was 2D Fourier-transformed. After extracting the low-frequency component of the image, the inverse Fourier transform was performed. Then, the blood vessel was detected by thresholding methods that is plugged in ImageJ as a function “AutoThreshold”. The algorithm when using this function was “MinError.” (3) Proximal ROIs with a high correlation coefficient of the signals. ROI pairs with ROI-to-ROI distances of 20 μm or less and partial correlation coefficients greater than 0.8 were removed for data analysis. More information on the calculation method of partial correlation coefficients is described within the section entitled “Topological analysis of functional networks.” See Table S4 and S5 for the number of ROIs used in the topological analysis.

### Movement analysis

Mouse body movements were measured by EMG of the neck muscles or by the gyro sensor fitted to the left HL. The EMG signals digitized at 20 kHz were bandpass filtered at 80-200 Hz using the MATLAB function “designfilt” and “filter.” As the designed filter was a finite impulse response filter, the delay caused by the bandpass filtering was constant. We obtained the filtered signal without delay by shifting the time calculated using the MATLAB function “grpdelay.” The periodic noise superimposed on the gyro signal was removed using a Savitzky–Golay filter. Then, we calculated normalized movement signals. The calculation process was identical for EMG signals and gyro sensor signals, as follows: (1) Decrease the sampling rate of the movement signals so that they matched imaging frame rates; (2) Subtract the DC component; (3) Integrate the absolute value of the filtered trace within each imaging frame interval; (4) Normalize the integrated trace between 0 and 1.

### Brain regions identification

Skull images including the cranial window of the head-fixed mouse was acquired from one-photon macroscopic imaging. The FOV was 16.6 × 14.0 mm^2^ or 13.3 × 13.3 mm^2^ and the image resolution was 1,280 × 1,080 or 1,024 × 1,024 pixels after 2x spatial binning (spatial resolution: ∼13 μm per pixel). An excitation light intensity from a blue LED source (LEX2-B-S, Brain Vision Inc.) and the gain and exposure time of the sCMOS camera (Zyla 5.5, Andor) was tuned so that all pixels were not saturated.

A median image of two-photon images for 500 frames was prepared, and the two-photon imaging region in the skull images was identified using the MATLAB function “imregtform.” Next, the Allen Mouse Common Coordinate Framework v3 (CCF) was fitted to the skull images in MATLAB. This fitting was performed using anatomical landmarks: bregma marked during surgery and rostral rhinal vein. In this fitting, the Allen CCF was rotated to fit the skull images, as the mouse was rotated in two-photon imaging so that the cranial window glass was positioned parallel to the focal plane of the objective lens.

### Calculation of decoding accuracy for movement

To identify neurons that carry information about movement, decoding analysis from ΔF/F signals to determine whether the animals move or not was performed. Twenty movement time points were chosen under the condition that the EMG signal increases significantly after more than 3 seconds of small EMG period. Here, the small EMG was defined as the normalized movement signal below 0.14 (see the section entitled “Movement analysis”). On the other hand, small EMG period that continues longer than 3 second is considered as no movement period. Twenty time points were randomly extracted from no movement period as representative points. The threshold of movement EMG signal is manually set to eliminate the individual difference. To discriminate the movement and no movement occurred at time *t*, ΔF/F signals at time *t*, *t* + 1, *t* + 2 were used. To extract the evoked activity by movement, the ΔF/F signals at time *t* − 1 were subtracted from the ΔF/F signal at time *t*, *t* + 1, *t* + 2.

Forty samples of movement and no movement data points (20 each) were classified using a support vector machine (SVM) with linear kernel. The SVM classifier was trained by using “fitcsvm” function in MATLAB. The correct classification rate was computed by 5-fold cross-validation. The method of selecting 20 samples of no movement data were changed 100 times and then, the correct rate was averaged over the 100 sets of the no movement data. The averaged correct rate obtained through this method was used as the decoding accuracy of each neuron.

### Topological analysis of functional networks

We estimated functional connectivity among the neurons based on Pearson’s correlation coefficient. As shown in our results (Fig. 5 and Fig. S5) and reported in previous studies(*28, 60*), a large number of cortical neurons demonstrate movement-related activity. Thus, in an effort to isolate resting-state associations among the neurons, we calculated partial correlations between each pair of neural activities by partialling out the contributions of correlations between the neural activities and the body movements measured with the gyro sensor. Normalized gyro sensor signals were shifted and averaged in a sliding window, in which a shifting time and a window size were determined so that the resulting partial correlations went to the minimum values. The above calculations were performed by using the “partialcorr” function in MATLAB.

Partial correlations were then used to construct undirected networks. We defined individual ROIs as nodes and partial correlations above the threshold as links that connect the correlated ROIs. To undergo the statistical assessment of a scale-free topology, the networks should be neither too sparse nor too dense(*21*); the mean number of links per node should be larger than 2 and smaller than the square root of the total number of nodes. Therefore, we set the threshold of the partial correlations at 0.4 to generate the links. A series of statistical tests to assess a scale-free topology was performed as previously described(*21, 24*) by using the python package available at https://github.com/adbroido/SFAnalysis.

Small-world topology was assessed by comparing the average shortest path lengths and the clustering coefficients of empirical networks with those of regular networks and random networks that contain the comparable number of nodes and links. Regular and random networks were generated using the “WattsStrogatz” function in MATLAB. Path lengths and clustering coefficients were calculated by using Gephi, an open source software for analyzing networks(*61*). For quantitative comparisons, we calculated two types of small-world metrics proposed in previous studies: the Small-world-ness(*25*) and the Small-world propensity(*26*).

### Histology

#### Tissue preparation

The mice were deeply anesthetized by intraperitoneal injection of urethane and perfused transcardially with 20 ml of HBSS (14025076, Life Technologies) supplemented with heparin (10 units/ml) and perfused with 4% formaldehyde in 0.1 M phosphate buffer (PB; pH 7.4), followed by postfixation in the same fixative for 16–20 h at 4 °C. After cryoprotection with 30% sucrose in PB, brain blocks were cut into 40-μm-thick sagittal sections or coronal sections on a freezing microtome (ROM-380, Yamato).

#### In situ hybridization

We carried out the hybridization procedure as reported in previous studies(*62*). The sense and anti-sense single-strand RNA probes for GAD67 (GenBank accession number: XM_133432.2) were synthesized with a digoxigenin (DIG) labeling kit (11277073910, Roche Diagnostics). Free-floating sections were hybridized for 16–20 h at 60 °C with a 1 µg/ml RNA sense or anti-sense probe in a hybridization buffer. After repeated washings and ribonuclease A (RNase A) treatment, the sections were incubated overnight with a mixture of 1:1000-diluted sheep anti-DIG-AP (11093274910, Roche Diagnostics) and 1:500-diluted rabbit anti-GFP antibody (598, Medical & Biological Laboratories Co., Ltd.) at 4 °C. The next day, the sections were incubated for 2 h at room temperature with Alexa Fluor 488 goat antibody (5 μg/ml) against rabbit IgG (A11034, LifeTechnologies), and finally reacted for 40 min with a 2-hydroxy-3-naphthoic acid-2’-phenylaniline phosphate Fluorescence Detection kit (1175888001, Roche Diagnostics). The sections were then placed on coverslips with CC/Mount (K002, Diagnostic Biosystems, Pleasanton).

#### Immunohistochemistry

Free-floating vibratome sections were incubated in a blocking solution (2% normal goat serum, 0.12% λ-carrageenan and 0.3% Triton X-100 in PBS) at room temperature for 1 h, followed by incubation with a 1:500-diluted primary rabbit anti-NeuN antibody (Neuronal marker, ABN78, Millipore) and a 1:500-diluted primary rabbit anti-Iba1 antibody (microglia marker, 019-19741, WAKO) overnight at 4°C. The next day, the sections were incubated for 2 h at room temperature with Alexa Fluor 647 goat antibody (5 µg/ml) against rabbit IgG (A21245, LifeTechnologies). The slices were mounted on glass slides and placed on coverslips with a Fluoromount/Plus anti-fading agent (K048, Diagnostic BioSystems).

#### Image acquisition and data analysis

The fluorescence-labeled sections were observed under a TCS SP8 confocal laser scanning microscope (Leica Microsystems) and an IX83P2 inverted microscope (Olympus). We could not detect higher signals than the background labeling using the sense probe. The number of L2/3 neurons that expressed G-CaMP7.09 and were positive for NeuN and GAD67 were manually counted using the “Cell Count” plug-in in ImageJ. The number of L2/3 neurons that expressed G-CaMP7.09 and were negative for NeuN was 37 (n = 13 brain sections, n = 3 mice. Note: we expected that not all neurons were stained with NeuN, and hereafter we did not use those cells for the estimation). The total count of NeuN positive neurons was 860 cells including 201 that were only NeuN positive cells and 659 that were NeuN and G-CaMP7.09 positive (merged) neurons. In situ hybridization to label inhibitory neurons showed that 99.5% of G-CaMP7.09 positive neurons were negative for GAD67, a marker of inhibitory neurons, in our experimental condition. It is known that 85-90% of cortical neurons are excitatory neurons(*63*). Together, 731-774 out of the 860 NeuN positive neurons were estimated to be excitatory neurons (85-90% of 860 cells). Due to the number of merged cells (NeuN and G-CaMP7.09) being 659, we expected that 85.1-90.2% out of all excitatory L2/3 neurons in those cortical areas were labeled in Fig. 4a-c (659 out of 731-774 cells).

### Point spread function

We imaged the 0.5 μm or 1 μm fluorescent beads embedded in agarose to estimate the point spread function (PSF). The beads had a maximum excitation wavelength of 505 nm and a maximum emission wavelength of 515 nm. The sample was sealed with a cover glass (0.13-0.17 mm in thickness, Matsunami Glass Ind.), and was set on a piezo stage (NANO-Z100-N, Mad City Labs Inc.) which moved along the Z-axis. The beads located ≤ 100 μm and 500 μm below the cover glass were imaged. Three-dimensional stack images of beads were collected along Z-axis by 1 μm steps. We acquired 22 or 60 layers through capturing 9 or 64 images for each layer. Following image capture, these images of each layer were averaged and were assumed as a representative image. The imaging parameters were as follows: 40 x 40 μm^2^ FOV; 2,048 x 2,048 pixels.

Due to a refractive index mismatch between the objective immersion medium (air) and the specimen medium (water), the stage displacement Δ*_stage_* dose matched the actual focal displacement Δ*_focus_*. As per a previous study(*64*), the relational equation between Δ*_stage_* and Δ*_focus_* was derived as follows:

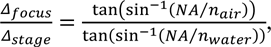

where *n_air_* = 1.00, and *n_water_* = 1.33. In this study, we substituted the 70% of the excitation NA into the NA in the above equation and finally obtained the correction as 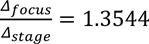.

We extracted the intensity profiles from the three-dimensional stack images. A Gaussian curve was fitted to the intensity profiles. The PSF was estimated as the full width at half-maximum (FWHM) of the fitted Gaussian curve.

### Region dependency of the signal-to-noise ratio

We recorded Ca^2+^ signals from the same neurons at the center, right, and top of the FOV (3.0 x 3.0 mm^2^). The recording procedure was previously described within the section entitled “In vivo two-photon imaging.” We used spontaneous activity to evaluate the signal-to-noise ratio. Firstly, we selected the neurons (n= 16) to be evaluated and positioned them at the center of the FOV. After recording the activity at the center of the FOV, the mouse was shifted by 1,406.25 μm (corresponding to 960 pixels) so that the selected neurons were positioned at the edges of the FOV (i.e., right or top of the FOV), then their activities were re-recorded. All the imaging parameters were the same at every region as follows: 3.0 mm x 3.0 mm FOV; 2,048 x 2,048 pixels; 7.5 frames/s; 130-μm depth below the pia.

After the recording, we identified each region where the selected neurons were located using the geometric transformation functions, “imregtform” and “imwarp,” in MATLAB. ROIs were manually drawn using the ImageJ “ROI Manager.” The size and shape of the ROIs were identical for each neuron at the center, right, and top of the FOV in order to eliminate changes in the signal-to-noise ratio caused by the ROI size and shape.

We extracted the fluorescence *F_ROI_* (t) of the selected neurons at the center, right, and top of the FOV. The ΔF/F was calculated according to procedures reported in the “Image analysis” section, with no neuropil correction performed. Importantly, this ΔF/F was not the denoised ΔF/F. We defined the signal as the ΔF/F below 0.5 Hz, and the noise as the ΔF/F above 0.5 Hz. We then calculated the signal-to-noise ratio using the “snr” function in MATLAB.

### Video

The ΔF/F movies in Video S1 and S2 were created in the following steps. (1) Brain motion of the raw imaging data (500 frames in total) was corrected using the ImageJ plugin “Image Stabilizer”. (2) A baseline image was calculated as a median projection of the 500 images obtained as a result in (1). (3) The ΔF/F images were calculated by dividing the brain motion corrected images by the baseline image. In this calculation, the images were extended from 16 bits to 32 bits in order to prevent rounding of values by division. (4) Noise superimposed on the ΔF/F images was removed using 3D median filter. Here, x radius and y radius were 0 and z radius was 3.0 in Video S1 and 12.0 in Video S2. (5) The bit depth was converted from 32 bits to 8bits and the contrast of the images was adjusted for visualization. In video 1, blood vessel was mask black. The detection of blood vessel was previously described within the section entitled “Image analysis”. (6) The images were output as an uncompressed avi file at a playback speed of 30 fps. The procedures in (1) – (6) were performed using ImageJ. We used VEGAS PRO 14 (Magix) to edit the video (one-photon CMOS image, raw movie and ΔF/F movie connection, and zooming in on the ΔF/F movies).

## Acknowledgments

We thank Y. Goda, H. Kamiguchi, M. Larkum, T. McHugh, S. Okabe and C. Yokoyama for comments and discussions on the manuscript; R. Endo, R. Kato, Y. Mizukami and K. Ueno for animal control; I. Oomoto for ImageJ program to detect blood vessels; A. Kamoshida for LabVIEW program to measure PSF; the staff of Nikon and Nikon Instruments for useful comments on the microscope; the staff of Hamamatsu Photonics K.K. for useful comments on the GaAsP PMTs; S. Itohara and K. Yamakawa for kindly providing VGAT-Cre mice; M. Nishiyama and Y. Kurokawa for comments on the videos; the RIKEN CBS CBS-Olympus Collaboration Center and Research Resources Division for supporting our anatomical experiments and imaging acquisition. This research was supported by the AMED-Brain/Minds Project under Grant Numbers JP19dm0207064 (issued to H.H.), JP16dm0207041 (issued to J.N.), and JP20dm0207001 (issued to A.M. and M.M.); by JST CREST (JPMJCR1864 and JPMJCR15E2) and JSPS KAKENHI (JP18H02713) through grants issued to M.O.; and by the Cooperative Study Program (237) of the National Institute for Physiological Sciences.

## Author contributions

M.M. designed the study. T.O., Y.K., M.M. and A.M. designed the microscope. M.H. developed the acquisition software. J.M. designed the GaAsP PMTs. K.O. performed the in vivo two-photon imaging. K.O., T.S., M.K. and Y.O. performed the viral injections and prepared the cranial windows. M. Odagawa, H.H. and K.O. performed the anatomical studies. M. Ohkura and J.N. created the G-CaMP7.09. C.M. and Y.O. performed the cloning for the AAVs. K.K. produced the AAVs. T.I., T.A. and K.O. performed cell detection using LCCD. M.I., Y.I., M. Oizumi and K.O. performed the data analysis. M.M., T.O., A.M., K.O., T.A., Y.I., and M. Oizumi prepared the manuscript. All authors contributed to the discussion of the experimental procedures, the results, and the manuscript.

## Supplementary Materials

**Figure S1.**
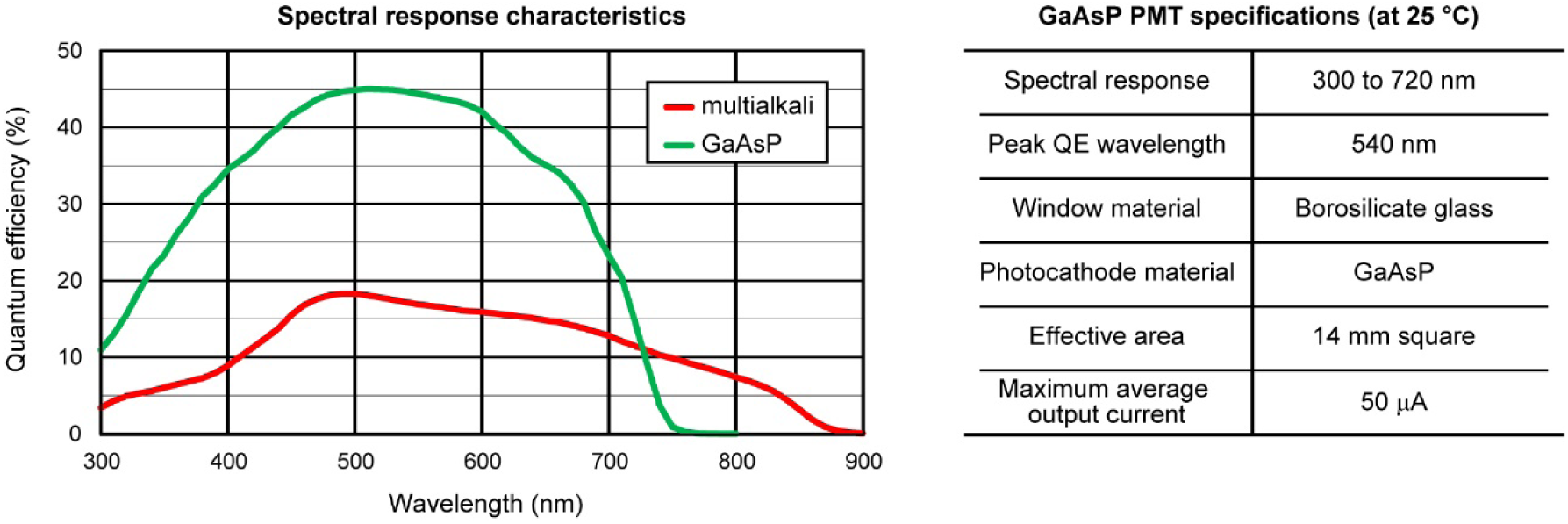
Spectral response characteristics and general specifications of the GaAsP PMT. Left: Quantum efficiencies (QEs) of the multialkali PMT (Hamamatsu Photonics K.K.) and the new GaAsP PMT. Right: General specifications of the GaAsP PMT.

**Figure S2.**
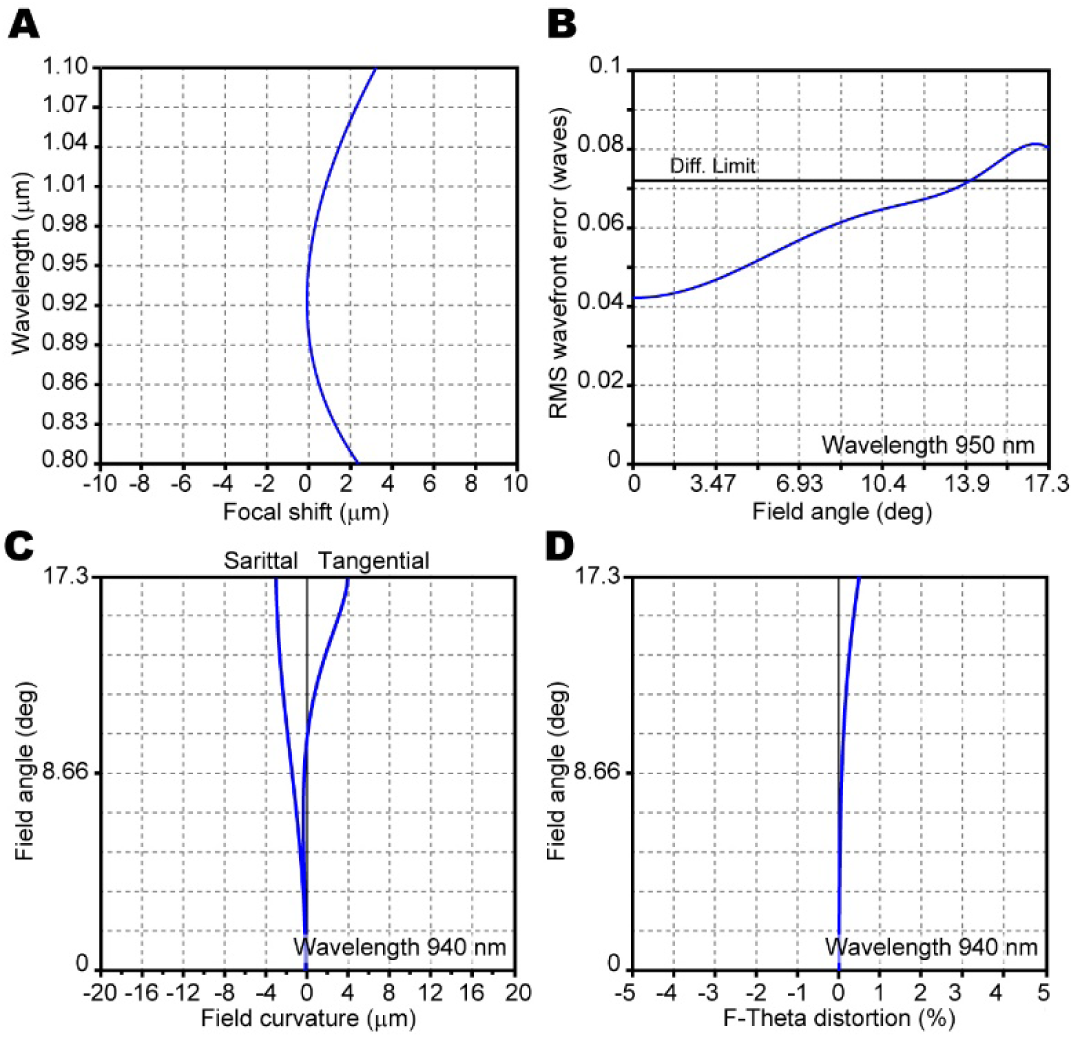
Optical performance of REGLOS. (**A-D**) Simulation of system performance. Chromatic aberration (**A**), root mean square (RMS) wavefront error (**B**), field curvature (**C**) and F-theta distortion (**D**) were simulated for a system including resonant and galvo mirrors, scan and tube lenses, a dichroic mirror and an objective with the actual distances between these components.

**Figure S3.**
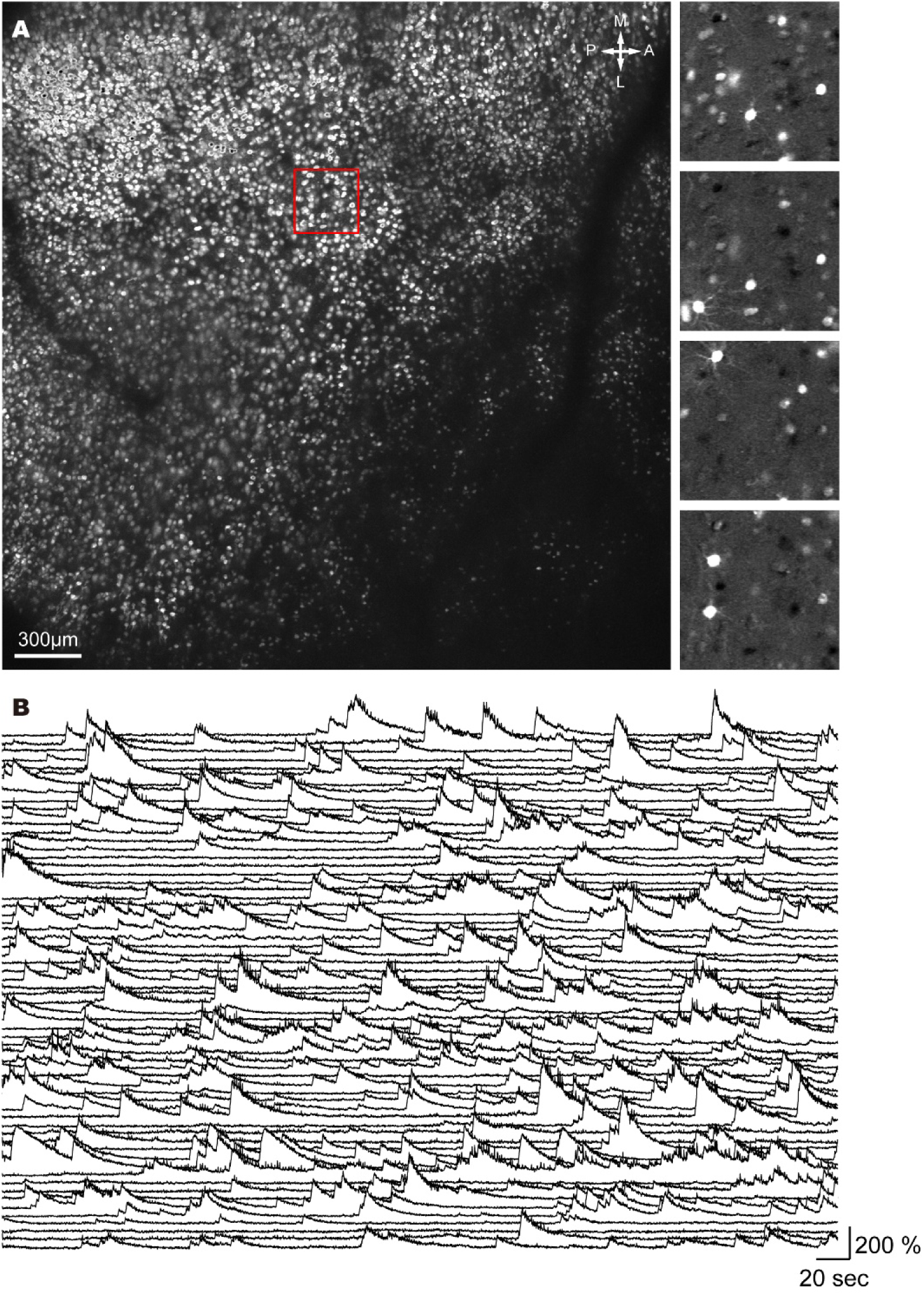
Ca^2+^ imaging from layer 5 neurons. (**A**) Left: A wide FOV (3 × 3 mm^2^) including layer 5 cortical neurons labeled with GCaMP6s (Rbp4-Cre mouse, AAV9-Syn-Flex-GCaMP6s, 302 mW laser power, 7.5 fps). Right: Examples of ΔF/F image from the area indicated by the red box in the left panel at different time points. (**B**) Randomly selected Ca^2+^ signals from 64 neurons shown on the right side in (**A**).

**Table S1.** Acquisition modes of FASHIO-2PM.

| Acquisition mode | Pixels | Pixel size ($\mu\text{m}$) | Area ($\text{mm}^2$) | Sampling rate (Hz) |
| --- | --- | --- | --- | --- |
| Mode 1 | 2048 x 2048 | 1.5 | 3 x 3 | 7.5 |
| Mode 2 | 512 x 512 | 5.9 | 3 x 3 | 30 |

**Table S2.** Lens specifications.

| Design of lens | Transmission<br>(nm) | Correction<br>(nm) | NA | WD (mm) | Image field<br>(mm<sup>2</sup>) | Focal length<br>(mm) | Aperture<br>diameter (mm) |
| --- | --- | --- | --- | --- | --- | --- | --- |
| Objective lens | 500-1000 | 800-1000 | 0.4/0.8 | 4.5 | 3 x 3 | 35 | 28/56 |
| Tube lens | 500-1000 | 500-1000 | 0.07 | 151 | 17.14 x 17.14 | 200 | 28 |
| Scan lens | 800-1000 | 800-1000 | 0.069 | 151 | 17.14 x 17.14 | 40 | 5.5 |

**Table S3.** Summary of imaging parameters.

| Figure | FOV | Pixels | Imaging speed | Imaging depth | PMT | Post-objective laser power |
| --- | --- | --- | --- | --- | --- | --- |
| Fig. 4E, G | 3.0 x 3.0 mm<sup>2</sup> | 2,048 x 2,048 | 7.5 frame/s | 165 µm | Multialkali | 176 mW |
| Fig. 4F | 0.75 x 0.75 mm<sup>2</sup> | 2,048 x 2,048 | 7.5 frame/s | 100 µm | Multialkali | 122 mW |
| Fig. 4H, I | 3.0 x 3.0 mm<sup>2</sup> | 2,048 x 2,048 | 7.5 frame/s | 130 µm | Multialkali | 148 mW |
| Fig. 5 & Video 1 | 3.0 x 3.0 mm<sup>2</sup> | 2,048 x 2,048 | 7.5 frame/s | 120 µm | Multialkali | 158 mW |
| Fig. 6 | 3.0 x 3.0 mm<sup>2</sup> | 2,048 x 2,048 | 7.5 frame/s | 110-130 µm | GaAsP (N=3)/<br>Multialkali | 62, 62 and 77mW<br>(G)/ |
| Video 2 | 3.0 x 3.0 mm<sup>2</sup> | 2,048 x 2,048 | 7.5 frame/s | 500 µm | GaAsP | 302 mW |

**Table S4.**
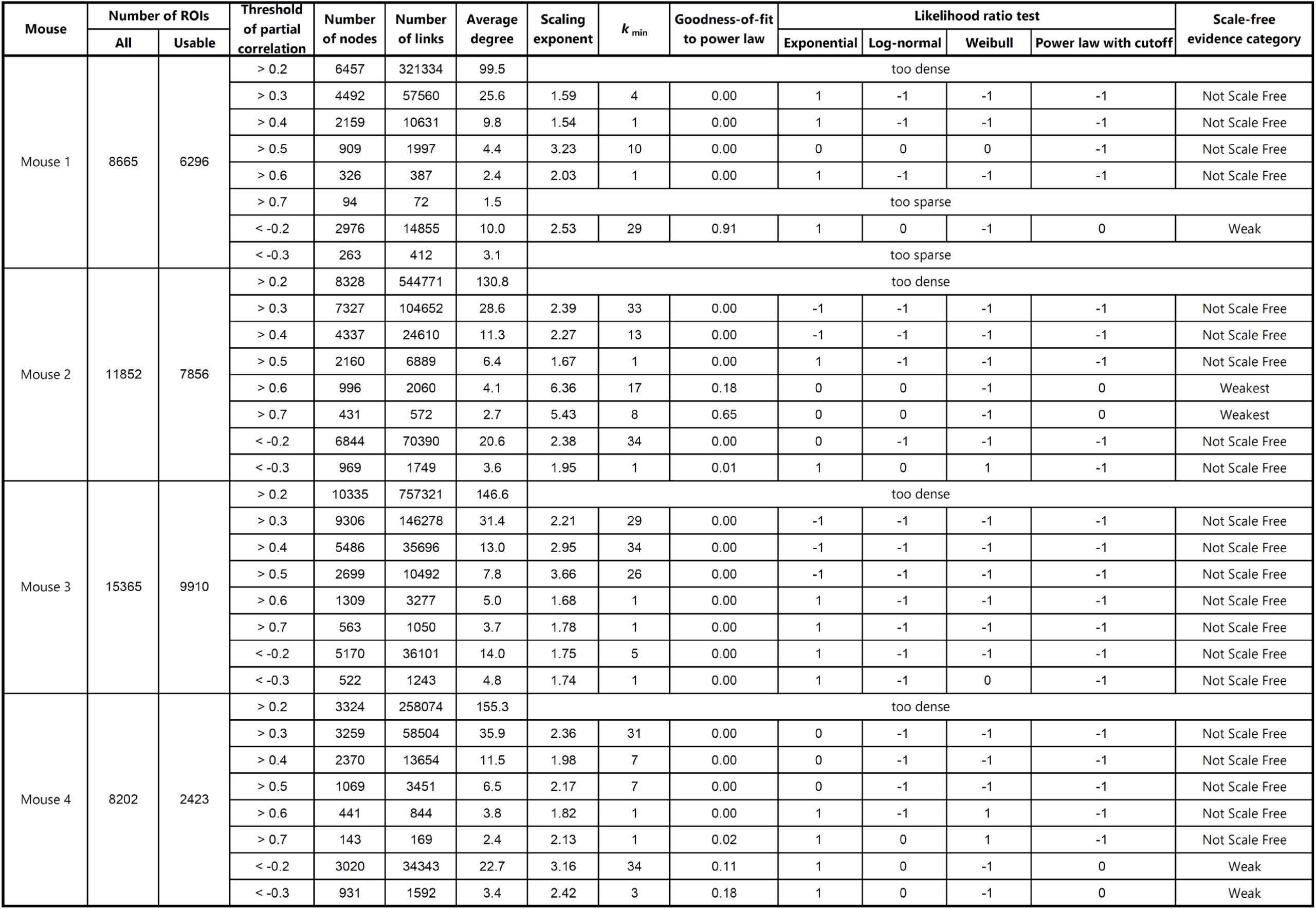
Statistical tests for scale-free properties. A series of statistical tests for scale-free properties were performed as previously described^6,7^ at different PCC criteria. All tests and comparisons were made only on the degrees *k* ≥ *k*_min_ in the upper tail, in which *k*_min_ was estimated so that a goodness-of-fit to power laws go to its maximum values. The sings of the likelihood ratio tests indicate which model is a better fit to the data: the power law (+1), the listed alternative distribution (−1), or neither (0).

**Table S5.**
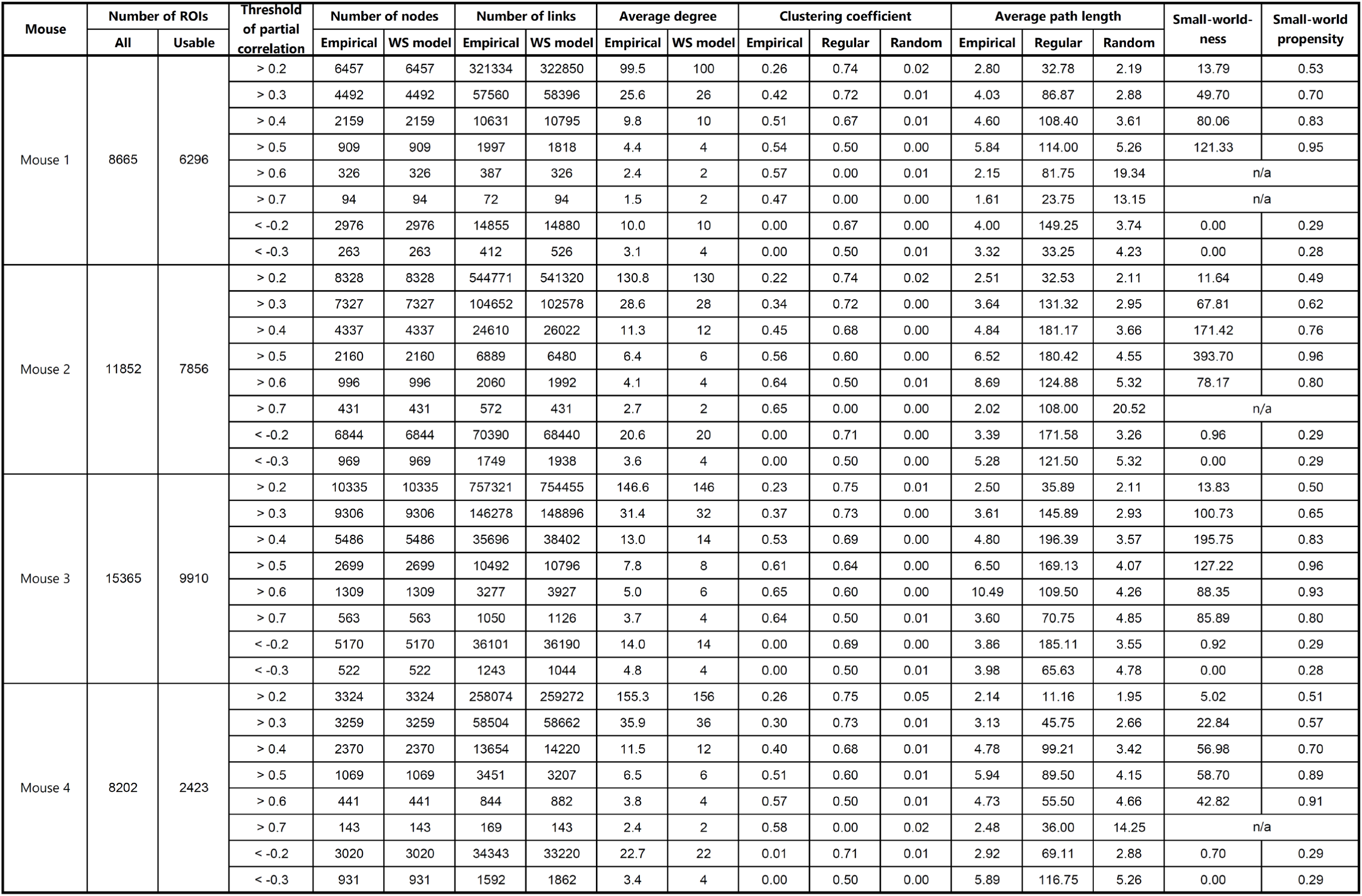
Network metrics for small-world properties. The average path lengths and clustering coefficients of cortical, regular, and random networks were calculated at different PCC criteria to assess the presence of small-world properties. Based on the standard Watts-Strogatz (WS) model, regular and random networks were generated by rewiring the links with probabilities of 0 and 1, respectively. Two types of small-world metrics, the small-world-ness and the small-world propensity, were calculated as previously described^29,30^.

## Captions of Supplementary Videos

**Video S1 Fast scanning high optical invariant two-photon microscopy for Ca^2+^ imaging from layer 2/3 neurons in an awake mouse.**

The raw imaging data are represented, followed by the ΔF/F representation. The data were recorded at 7.5Hz (without averaging frames, 158 mW laser power, see Supplementary Table 3 for other parameters) from 3 x 3 mm^2^ FOV, and are play back at 4x real-speed. As pre-processing the movie, we corrected for brain motion artifacts, and adjusted the contrast to highlight the details (Methods).

**Video S2 Fast scanning high optical invariant two-photon microscopy for Ca^2+^ imaging from layer 5 neurons in an awake mouse.**

The ΔF/F representation. The data were recorded at 7.5Hz (without averaging frames, 302 mW laser power, see Supplementary Table 3 for other parameters) from 3 x 3 mm^2^ FOV, and are play back at 4x real-speed. As pre-processing the movie, we corrected for brain motion artifacts, and adjusted the contrast to highlight the details (Methods).

